# Type VI Secretion Promotes Microbial Antagonism and Contributes to Pathogenesis by AHPND-causing *Vibrio parahaemolyticus*

**DOI:** 10.1101/2024.06.03.597196

**Authors:** Damian Cavanagh, Karina L. Martinez Lopez, Jack Headrick, Katie Sprinkel, Monique L. van Hoek, Brett Froelich

## Abstract

The Type VI Secretion System (T6SS) is a contractile injection apparatus which delivers effector toxins in a contact-dependent manner, and has drawn considerable attention as a determinant of pathogenicity and environmental fitness in diverse bacterial taxa. Past reports have noted that many strains of *Vibrio parahaemolyticus* which cause Acute Hepatopancreatic Necrosis Disease (AHPND) in shrimp harbor two functional T6SSs (T6SS1 and T6SS2), leading to speculation that these systems may contribute to virulence. Here, we evaluate the involvement of Type VI Secretion (T6S) as a pathogenicity factor during shrimp infection by two geographically distinct isolates of AHPND-causing *V. parahaemolyticus* (Vp_AHPND_). Combining in vitro contact-killing assays with comparative genomic analysis, we demonstrate that these strains — 13-306/D4 (Mexico) and 12-297/B (Vietnam) — leverage overlapping yet distinct effector arsenals to antagonize bacterial and fungal prey under warm, high-salinity conditions. Moreover, our in silico analyses reveal a predicted T6SS effector/immunity (E/I) module encoded in proximity to the *pirAB*-Tn903 composite transposon on pVA1-type plasmids. We demonstrate that acquisition of a plasmid expressing this module allows the non-AHPND isolate *V. parahaemolyticus* BB22OP to antagonize its parental strain in a T6SS1-dependent fashion. In turn, prey equipped with a plasmid expressing only the immunity gene gain protection against BB22OP attackers armed with the full E/I module. Employing an immersion bioassay, we show that both D4 and 297B require T6S functionality for full virulence against whiteleg shrimp (*Litopenaeus vannamei*) postlarvae under warm, marine-like conditions, and provide preliminary evidence that T6SS-dependent interactions with host-associated bacteria may contribute to virulence. Our findings offer empirical evidence supporting the involvement of T6S in AHPND pathogenesis under conditions relevant to commercial aquaculture, inviting further studies to decipher the role/s of T6S as a virulence determinant.

## Introduction

Shrimp aquaculture constitutes one of the largest and fastest-growing sectors of food production throughout the world and serves as one of the major sources for high-protein seafood in South America, China, Thailand, Malaysia, and Vietnam. Given the scale of the shrimp aquaculture industry, production losses caused by infectious diseases such as Acute Hepatopancreatic Necrosis Disease (AHPND) can prove economically devastating [1, 2].

Outbreaks of AHPND (formerly called Early Mortality Syndrome) often occur within the first 20-35 days of shrimp grow-out during the postlarval stage, and are noted for causing mortality rates approaching 100% in less than one week following the appearance of clinical signs [1–3]. Multiple species of commercially farmed shrimp are susceptible to AHPND, including *Penaeus monodon* (black tiger shrimp) and *Litopenaeus vannamei* (whiteleg shrimp), the latter of which accounted for over 53% of global crustacean production totals in 2016 [4]. Cumulative financial losses attributable to AHPND from the years 2010 to 2016 have been estimated at over $44 billion USD, and the list of territories reporting outbreaks of AHPND has expanded in recent years [1, 5–8]. Furthermore, communities that lean heavily on shrimp aquaculture for income and food production are severely impacted by mass mortality events caused by AHPND.

The economic harm posed by AHPND has necessitated investigations to identify the etiologic agents and key virulence factors responsible for this illness. Past works have demonstrated that AHPND is mainly propagated by a subset of *Vibrio parahaemolyticus* strains (Vp_AHPND_), although isolates of related species such as *Vibrio harveyi* and *Vibrio owensii* have subsequently been confirmed as causative agents as well [9–15]. Regardless of species, AHPND-causing strains are distinguished by possession of self-transmissible virulence plasmids which express a pesticidal *Photorhabdus* insect-related (Pir) family binary toxin [5, 9, 10, 16–19]. These plasmids, represented by pVA1, harbor the *pirA^vp^*/*pirB^vp^* toxin genes within an unstable composite transposon [13–16]. In addition to the *pirAB*-Tn903 mobile element, pVA1-type plasmids encode genes to facilitate conjugative transfer, and are maintained by a hok/gef family post-segregational killing (PSK) system [20–22].

The PirA/B^vp^ binary toxins are established as the major virulence determinants of AHPND, and are required to induce necrosis of the shrimp hepatopancreas [5, 9, 12]. However, Pir toxin production alone is insufficient to explain successful colonization and persistence within the shrimp host.

Consequently, it has been speculated that other factors are involved in host adaptation and disease progression. In particular, Type VI Secretion (T6S) is frequently cited as a possible virulence determinant in the context of shrimp infection by Vp_AHPND_ [23–27].

The Type VI Secretion System (T6SS) is a versatile, spear-like molecular complex which is broadly employed by gram-negative bacteria to convey effector toxins and other protein cargo [28, 29]. The T6SS apparatus consists of a central tube of stacking Hemolysin-coregulated protein (Hcp) monomers encompassed by an outer sheath and terminating in a trimeric Valine-glycine repeat protein G (VgrG) spike **(Fig. 1)**. The VgrG spike is capped by a Proline-Alanine-Alanine-aRginine (PAAR) domain-containing protein that serves to sharpen the tip of the “spear” for more effective puncturing. When fired, the central tube of the T6SS is propelled forward by contraction of the sheath, which is anchored to the cell envelope by a baseplate complex [29]. Delivery of substrate molecules by the T6SS is accomplished through various means, and effector proteins are typically loaded within the lumen of the Hcp tube or else attached to the VgrG-PAAR spike complex by a delivery domain [30].

**Fig. 1.**
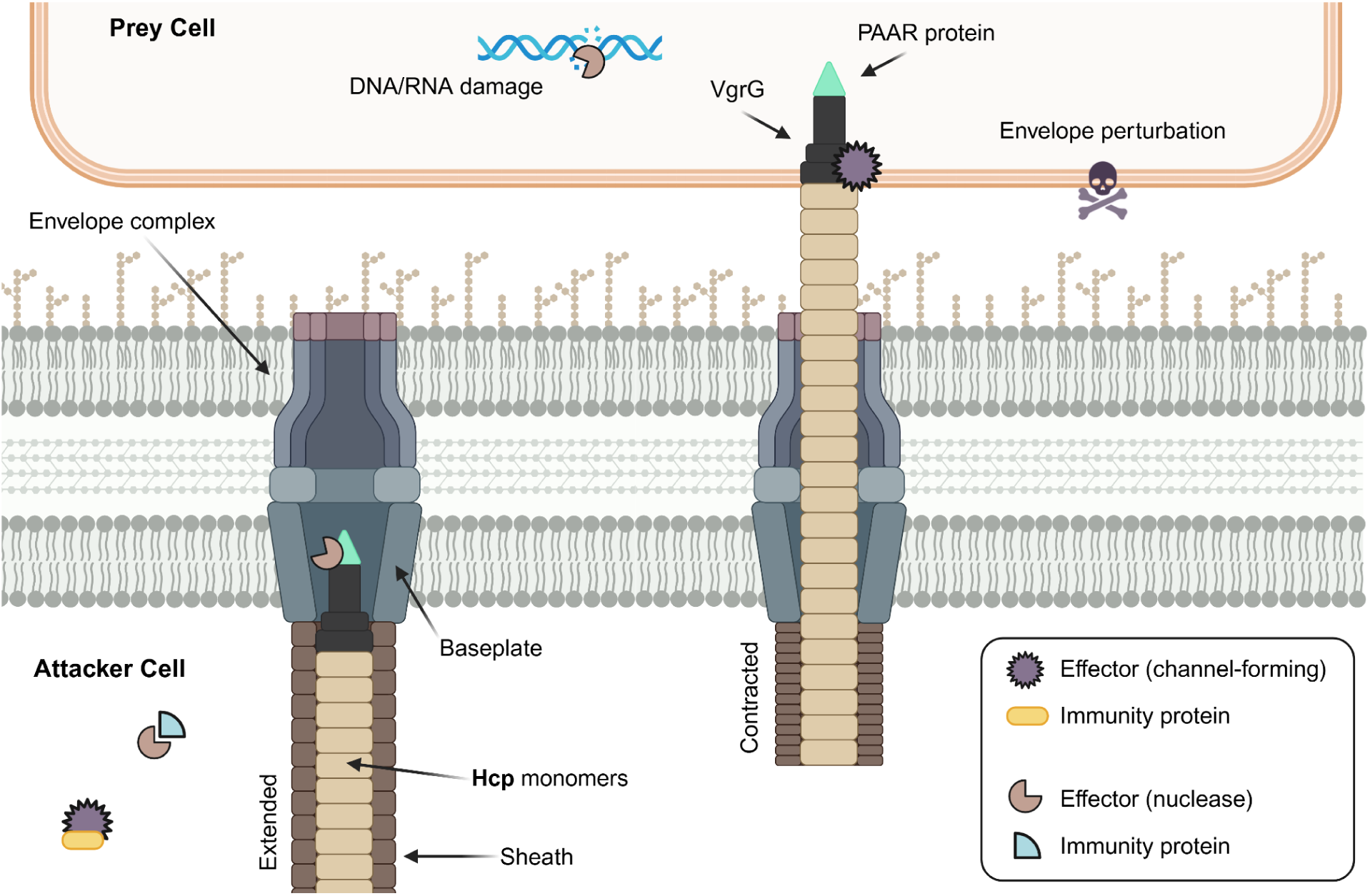
Overview of the Type VI Secretion System (T6SS). The central apparatus of the T6SS nanomachine consists of a tube made up of stacking Hcp monomers which terminate in a VgrG-PAAR spike complex. These components are surrounded by an outer sheath and anchored to the cell envelope by a baseplate and membrane complex. Contraction (firing) of the device enables a fusillade of effectors to be delivered into a neighboring cell.

A wide array of T6SS effector molecules have been reported in the literature, with many of these functioning as toxins that inhibit or kill the recipient [30–33]. Effectors are often categorized on the basis of target specificity (e.g. as antibacterial, anti-host, or antifungal effectors) and can also be grouped according to mechanism (e.g. pore-formation, metabolic perturbation, nuclease activity) [30, 34]. Importantly, antibacterial effector genes almost always neighbor an immunity gene which expresses an “antidote” protein to mitigate self-intoxication and kin-killing [30].

Type VI Secretion (T6S) has been implicated as a factor in the competitive fitness and virulence of a wide range of bacterial taxa, many of which belong to clinically significant genera such as *Yersinia*, *Pseudomonas*, and *Vibrio* [28, 35–38]. Some species that utilize T6S harbor more than one functional T6SS, each of which may be regulated independently and employed in different scenarios. Further, species possessing more than one T6SS may deliver a different payload of protein effectors with each [35].

Past reports have noted that, in addition to the canonical T6SS2 cluster found in nearly all strains of *V. parahaemolyticus*, most Vp_AHPND_ isolates possess a T6SS1 core cluster [23–27]. Studies of T6S functionality in *V. parahaemolyticus* have demonstrated the antibacterial activities of T6SS1 and T6SS2 and characterized the regulatory networks to which these systems are subject [23, 39–44]. Within this framework, it has been speculated that one or both of these T6SSs confer a fitness advantage that allows Vp_AHPND_ to overcome and displace the native shrimp microbiota, thereby facilitating colonization of the host [23, 24, 26, 27, 45, 46]. However, direct investigations of T6SS1 and T6SS2 as factors influencing AHPND progression are lacking. In this study, we empirically assess the extent to which these systems impact post-infection mortality in *L. vannamei* postlarvae challenged with the representative Vp_AHPND_ strains 13-306/D4 (D4) and 12-297/B (297B).

## Materials and methods

### Bacterial strains and media used

The AHPND-causing *V. parahaemolyticus* strains 13-306/D4 and 12-297/B were kindly provided by Kim Orth (University of Texas Southwestern Medical Center), while the non-AHPND isolate BB22OP was obtained as a gift from Dor Salomon (Tel Aviv University). *V. parahaemolyticus* strains were maintained on marine minimal medium (MMM) agar plates (1.5% [wt/vol] agar, 2% [wt/vol] NaCl, 0.4% [wt/vol] galactose, 5 mM MgSO_4_, 7 mM K_2_SO_4_, 77 mM K_2_HPO_4_, 35 mM KH_2_PO_4_, 2 mM NH_4_Cl), and routinely cultured at 30°C with continuous shaking in marine Luria-Bertani (MLB; LB adjusted to 3% [wt/vol] NaCl) or in tryptic soy broth supplemented with 2% [wt/vol] NaCl (TSB+) as indicated [23, 47].

*E. coli* strain S17-1 (λ-pir) was used for maintenance and conjugation of pDM4 and derivative plasmids. *E. coli* DH5α and commercial derivatives were used for cloning and plasmid maintenance, and as a prey strain in competition experiments. Ectopic expression from arabinose-inducible vectors was carried out in *E. coli* BL21-AI. All *E. coli* strains were cultured at 37°C in LB broth or on LB agar plates, supplemented with chloramphenicol (20 μg/mL) and/or kanamycin (50 μg/mL) when necessary for plasmid maintenance. The fungal strain *C. albicans* ATCC 90028 was routinely cultured in yeast extract peptone dextrose (YPD) broth or on YPD agar plates supplemented with chloramphenicol (20 μg/mL). A list of bacterial strains used in this study is provided in **Supplementary Table S1**.

### Construction of in-frame deletion mutants

Mutants of *V. parahaemolyticus* harboring clean, in-frame deletions of target genes were generated via allelic exchange using the CmR OriR6K suicide plasmid pDM4 as previously reported [23, 48, 49]. In brief, 1-kb regions directly upstream and downstream of the target gene were amplified from a genomic DNA template and inserted via Gibson Assembly into a pDM4 plasmid linearized with SacI-HF. The resulting constructs were first transformed into *E. coli* S17-1 (λ-pir) and then subsequently conjugated into *V. parahaemolyticus*. Transconjugants were selected for on MMM supplemented with chloramphenicol (10 μg/mL), screened for sucrose sensitivity, and then patched onto plain MMM agar for outgrowth. Counterselection was performed on sucrose-containing MMM plates, and putative secondary recombinants were checked for chloramphenicol sensitivity. Allelic exchange mutants harboring the target deletions were identified by PCR **(Supplementary Fig. S1a** and **S1b)**. To confirm retention of the *pirA* and *pirB* genes, mutant derivatives of D4 and 297B were screened using the duplex PCR protocol described by Han et al. (2015) [11]. Plasmids and primer sequences used in this work are listed in **Supplementary Tables S2** and **S3**, respectively.

### Growth experiments

Growth curve experiments were carried out on a plate reader as previously described, with minor modifications [47]. Briefly, starter cultures of *V. parahaemolyticus* wild type and mutant strains were grown in MLB overnight at 30°C with continuous shaking. The following day, cultures were washed with PBS and normalized to an OD_600_ of 0.1 in fresh MLB, then aliquoted in quadruplicate (200 μL per well) into a 96-well plate. Plates were incubated in a BioTek Cytation 5 microplate reader (Agilent, Santa Clara, CA) at 30°C with continuous shaking. Growth was measured as OD_600_ every hour over the course of 8 hours. Experiments were repeated three times.

### Plasmid manipulations

For arabinose-inducible, glucose-repressible expression of WP_025789713.1, the CDS for ACMX29_RS24340 (without a stop codon), was amplified from *V. parahaemolyticus* 13-306/D4 and cloned into pPER5 in-frame with an N-terminal pelB leader sequence and a C-terminal myc-His tag by means of Gibson Assembly [42, 50]. To express WP_025789712.1, the CDS for ACMX29_RS24335 was likewise amplified and assembled into the multiple cloning site (MCS) of pBAD33.1F in-frame with a C-terminal FLAG tag sequence.

Plasmids used for ectopic expression in *V. parahaemolyticus* were constructed using the mobilizable, broad host range (RSF1010 family) plasmid pJRD215 as a backbone [51]. Primers suitable for Gibson Assembly into the ApaI site of pJRD215 were used to amplify regions spanning from 27-bp upstream of the *araC* stop codon to 30-bp downstream of the *rrnB* T2 terminator in pBAD33.1F and derivatives. To generate a broad-host plasmid expressing the WP_025789713.1/WP_025789712.1 module, the CDS for ACMX29_RS24340-35 was amplified from D4 genomic DNA and inserted into the MCS of pBAD33.1F, in-frame with a C-terminal FLAG tag. The resulting intermediate plasmid was used to amplify a single fragment encompassing *araC*, the *araBAD* promoter, the ACMX29_RS24340-35 CDS, and the *rrnB* terminator sequences, which were then introduced into an ApaI-digested pJRD215 to yield pJRD_ara-Module. An empty MCS control plasmid (pJRD_ara-Empty) was generated in similar fashion using a fragment amplified from the original pBAD33.1F.

For prey selection and expression of the WP_025789712.1 immunity protein in BB22OP, derivatives of pJRD215 containing a tetracycline resistance marker (pJRD215-Tc) were used. First, the KmR cassette of pJRD215 was excised by digesting with XhoI and BmtI-HF, and subsequently replaced by a cassette harboring TetA and TetR derived from the RK2 family plasmid pCM62 [52]. Next, primers suitable for Gibson Assembly into the BamHI site of pJRD215-Tc were used to amplify regions spanning from 27-bp upstream of the *araC* stop codon to 30-bp downstream of the *rrnB* T2 terminator in pBAD33.1F and derivatives. Fragments amplified from pBAD33.1F_WP_025789712.1 and an empty pBAD33.1F were then assembled into BamHI digested pJRD215-Tc to generate pJRD-Tc_ara-Imm and the control plasmid pJRD-Tc_ara-Empty, respectively.

Assembled plasmids were introduced into S17-1 (λ-pir) and mobilized into BB22OP wild type and mutant recipients via conjugation. Transconjugants were obtained on MMM agar plates supplemented with kanamycin (250 μg/mL) or tetracycline (10 μg/mL) as appropriate, and then PCR screened to confirm the presence of inserts. Plasmid constructs were additionally verified by whole-plasmid sequencing (Oxford Nanopore, R10.4.1). Further details on plasmid construction are made available in the **supplementary information** for this manuscript.

### Bacterial and fungal competition assays

Contact-killing experiments were performed as described previously [23]. Overnight cultures grown in MLB (*V. parahaemolyticus*) or LB (*E. coli*) were pelleted, washed twice with phosphate-buffered saline (PBS), and normalized to an optical density at 600 nm (OD_600_) of 0.5 before being mixed at a 4:1 ratio (attacker-to-prey). Mixed cells were then spotted (25 μL) in triplicate onto MLB agar plates and incubated for 4 hours at 30°C. Prey concentrations (in CFU/mL) at the start of the assay (T0) were determined by spotting 10-fold serial dilutions of mixed cells onto LB agar containing 20 μg/mL chloramphenicol for selection of *E. coli* harboring an empty pBAD33.1 plasmid. At the end of the assay (T4), competition spots were harvested, and surviving prey were enumerated by spotting 10-fold serial dilutions as above.

Competition assays against fungal prey were carried out as described previously, with some modifications [53]. Attacker and prey strains grown overnight in MLB (*V. parahaemolyticus*) or YPD (*C. albicans*) were adjusted to an OD_600_ of 0.5 as above, then mixed at a 10:1 ratio (attacker-to-prey) and spotted (25 μL) onto TSA+ in triplicate. Following 8 hours of co-incubation at 30°C, competition spots were harvested and resuspended in PBS. Prey enumeration was carried out at the start and end of the assay by spotting serial 10-fold dilutions onto YPD agar supplemented with chloramphenicol.

Competitions between plasmid-bearing strains of BB22OP were performed as follows. Attacker and prey strains grown overnight in MLB were pelleted, washed twice with PBS, and normalized to an OD_600_ of 0.5. Cells were then mixed at a 10:1 ratio (attacker-to-prey) and spotted (25 μL) in triplicate onto MLB agar plates (pH 8) supplemented with 0.1% or 0.02% arabinose as indicated (to induce expression driven by the *araBAD* promoter). Competition spots were incubated for 4 hours at 30°C, after which time they were harvested. Prey enumeration was carried out at the start (T0) and end (T4) of the assay by spotting 10-fold serial dilutions onto MMM agar plates supplemented with tetracycline (10 μg/mL).

Competition experiments were performed three times (*n* = 3) in technical triplicate, and statistical significance between samples at T4 was determined using two-way ANOVA (matched by biological replicate) with either Tukey’s test for multiple comparisons or Fisher’s LSD test as appropriate.

### Sequence analysis and effector prediction

An updated genome assembly (<u>GCF_050888945.2</u>) for *Vibrio parahaemolyticus* 13-306/D4 was posted to the NCBI database with RefSeq accessions <u>NZ_CP180435.1</u> (Chromosome 1), <u>NZ_CP180436.1</u> (Chromosome 2), and <u>NZ_CP180437.2</u> (Plasmid pVA). The NCBI BLAST suite was employed to survey the genomes of *V. parahaemolyticus* 13-306/D4 and 12-297/B (<u>GCF_002114585.1</u>) for homologs of previously reported effectors [54]. Amino acid FASTA files for both genomes were obtained from The Rapid Annotations using Subsystems Technology (RAST) server (https://rast.nmpdr.org/) and used as queries against a curated list of T6SS effector proteins from the literature [55]. Protein accession and locus information for predicted effectors was gathered from the NCBI database, and is presented along with alignment results in **Supplementary Tables S4** and **S5**.

Structure prediction and domain analyses were carried out using InterPro release 107.0 (https://www.ebi.ac.uk/interpro/) and the MPI Bioinformatics Toolkit (https://toolkit.tuebingen.mpg.de/) [56, 57]. Default parameters were used for all software unless otherwise noted.

### Bacterial toxicity assays

To assess toxicity resulting from expression of WP_025789713.1, either alone or co-expressed with WP_025789712.1, *E. coli* BL21-AI were co-transformed with combinations of pPER5_WP_025789713.1, pBAD33.1F_WP_025789712.1, or the respective empty vector controls.

Fresh co-transformants were inoculated into LB supplemented with 0.4% [wt/vol] glucose, 50 μg/mL kanamycin, and 20 μg/mL chloramphenicol and grown overnight at 37°C with constant shaking. The following day, cultures were washed twice and normalized to an OD_600_ of 1.0 in sterile PBS. Serial 10-fold dilutions made in PBS were then spotted onto LB agar containing 50 μg/mL kanamycin, 20 μg/mL chloramphenicol, and either 0.4% [wt/vol] glucose or 0.02% [wt/vol] arabinose for repression or induction of expression, respectively. Plates were inspected visually after a 24 hour incubation at 37°C.

### L. vannamei maintenance

Disease-free *L. vannamei* postlarvae (PL) derived from a Specific-Pathogen-Free (SPF) broodstock were supplied by Miami Aquaculture (Boynton Beach, Florida, USA), and maintained in an aquarium tank filled with approximately 25 L of Artificial Seawater (ASW; 30 grams Instant Ocean® per liter of deionized water) with filtration (Tetra Whisper EX30 Power Filter) and aeration with attached airstones. Maintenance tank water was kept at a temperature of approximately 30°C using an Ultra-Slim Submersible Aquarium Heater (Fluval, Montreal, Canada). Tank water was maintained at a pH of 7.6 ± 0.4 and salinity of approximately 30 parts per thousand (ppt). Lighting was provided on a 12 hour schedule, and tank water was refreshed on a biweekly basis. PL were fed an estimated 10% body weight per day of PL Raceway Plus feed (Zeigler, Gardners, PA).

### L. vannamei infections

Infection experiments were carried out using a modified version of the immersion bioassay described by Tran et al. (2013) [9]. Briefly, isolated colonies of *V. parahaemolyticus* maintained on MMM agar plates were inoculated into TSB+ and grown overnight at 30°C with continuous shaking. The following day, *L. vannamei* ranging from PL24 to PL32 (average weight 0.29 ± 0.11 g) were allocated into beakers containing sterile ASW (2 beakers for each treatment group, with 6 shrimp per beaker), and inoculated with either sterile PBS or a sufficient volume of the indicated *V. parahaemolyticus* strain to achieve a concentration of approximately 1×10^8^ CFU/mL. After a 15 minute immersion at 30°C, PL were transferred into foil-covered experiment containers filled with 400 mL of sterile ASW and reinoculated with either PBS or a volume of bacterial suspension corresponding to a final density of approximately 1×10^6^ CFU/mL.

Experiment containers were maintained at a temperature of 30°C and salinity of 30 ppt, with aeration provided by sterile airstones. Lighting was provided on a 12 hour cycle, and PL were fed approximately 10% of their estimated body weight once per day. Survival rates were determined at 12, 24, and 48 hours post-infection. Individuals that displayed signs of atrophy and did not move for 5 minutes during observation were defined as nonviable.

For challenge experiments using antibiotic-treated *L. vannamei*, PL were incubated for 24 hours at 30°C in sterile ASW supplemented with colistin (10 µg/mL), tetracycline (10 µg/mL), ampicillin (100 µg/mL), and vancomycin (50 µg/mL). Parameters used for the antibiotic pre-treatment step were kept consistent with those maintained throughout the remainder of the challenge (i.e., 12 hour lighting cycles, aeration provided by sterile airstones, and a commercial feed diet corresponding to ∼10% body weight once daily). To minimize residual antibiotic carryover prior to bacterial challenge, antibiotic-supplemented ASW was replaced with fresh, sterile antibiotic-free ASW before proceeding with inoculation as described above.

Infection bioassays were carried out in 2 replicate containers for each treatment group, with 6 shrimp per container. Two independent experiments were performed, using a fresh cohort of shrimp for each. Survival trajectories were checked for consistency across duplicate containers and independent trials before pooling data for presentation. Kaplan-Meier survival curves were generated in GraphPad Prism, and statistical differences between groups were evaluated by log-rank test. Because PL within the same container shared water and simultaneous bacterial exposure, containers (*n* = 4 total containers per treatment group) rather than individual shrimp (*N* = 24 total animals per treatment group) may be regarded as a more conservative unit of replication, which is noted as a limitation of the statistical framework applied here.

### Midgut sample collection

Experiment containers filled with 400 mL of sterile, antibiotic-treated or untreated ASW were stocked with *L. vannamei* (6 shrimp per container) and incubated at 30°C for 24 hours with aeration provided by sterile airstones. Midgut samples (hepatopancreas and intestines) were collected from 3 shrimp per container (3 antibiotic-treated and 3 untreated shrimp per trial) following a modified procedure based on Soto-Rodriguez et al. (2015) [58]. Briefly, *L. vannamei* were euthanized by placing their containers on ice, then rinsed with 70% ethanol followed by sterile ASW. Midguts were aseptically dissected using a sterile blade and placed into microcentrifuge tubes for weighing, then homogenized individually in 100 µL of sterile ASW for every 10 mg of tissue. Serial 10-fold dilutions of the resulting homogenates were made in sterile ASW and plated on MLB agar. Plates were incubated at 30°C for 24 hours and then inspected for CFU determinations. Two independent experimental trials were performed.

### Antibiotic susceptibility testing

Predictions of the antibiotic resistance profile of *V. parahaemolyticus* 13-306/D4 were made based on literature reports of resistance determinants present in the D4 genome, including the *vpa0477*-like, *tetB*, and *dgkA-eptA* loci conferring resistance to ampicillin, tetracycline, and colistin respectively **(Supplementary Table S7)** [59–61]. Drug resistance phenotypes were then determined experimentally using a modified agar dilution method **(Supplementary Fig. S3)** [62]. Overnight cultures of wild type and mutant strains of D4 were grown in cation-adjusted Mueller-Hinton (CAMH) broth supplemented with 1% NaCl at 30°C with continuous shaking [63, 64]. The next day, cultures were normalized to a concentration of approximately 1.5×10^8^ CFU/mL. Serial 10-fold dilutions were then spotted onto MMM and CAMH + 1% NaCl agar plates supplemented with 10 µg/mL tetracycline (Tet10), 10 µg/mL colistin (Col10), 100 µg/mL ampicillin (Amp100), and 50 µg/mL vancomycin (Van50), or onto unsupplemented MMM. Plates were briefly allowed to dry, and incubated at 30°C for 24h before inspection.

### RNA extractions and RT-qPCR

Culture samples used for RNA extraction and isolation were prepared as follows. Individual colonies of *V. parahaemolyticus* 13-306/D4 wild type and mutant strains were inoculated from MMM agar plates into TSB+ and grown overnight at 30°C with constant shaking. The next day, cultures were diluted 1:100 in fresh TSB+, and grown for an additional 4 hours at 30°C. Extraction and isolation of RNA was performed following the TRIzol reagent protocol (Invitrogen). To minimize DNA contamination, samples were treated with TURBO DNase (Ambion).

The expression levels of *pirA^vp^*, *pirB^vp^*, and an internal control gene *gyrB* (DNA gyrase B, a housekeeping gene), were quantified using a Power SYBR Green RNA-to-CT 1-Step Kit (Applied Biosystems) following the manufacturer’s protocol. Primer pairs used for each target sequence have been described previously, and are listed in the **supplementary information** [65]. Reactions were carried out in a MyiQ Single-Color Real-Time PCR Detection System (Bio-Rad), using the following program: 48°C for 30 min, 95°C for 10 min, followed by 40 cycles of 95°C for 15 sec and 60°C for 1 min. A melt curve was also performed according to the manufacturer’s protocol.

Data from RT-qPCR experiments were analyzed by the comparative C_q_ method. For each biological replicate, ΔC_q_ was calculated by subtracting the C_q_ value of the internal reference gene (*gyrB*) from that of the target gene (*pirA^vp^* or *pirB^vp^*), such that lower (more negative) ΔC_q_ values correspond to greater target transcript abundance. The ΔΔC_q_ values were calculated as the difference between the ΔC_q_ of each mutant strain and that of wild type D4, and relative expression was expressed as 2^−ΔΔCq^, assuming an amplification efficiency of 2. Under this convention a negative ΔΔC_q_ denotes increased target abundance in the mutant relative to wild type, and a relative expression value greater than 1 denotes upregulation.

Separate one-way analyses of variance were performed on the ΔC_q_ values for *pirA^vp^* and *pirB^vp^* followed by Dunnett’s multiple comparisons test against wild type D4. Analyses were conducted on the ΔC_q_ scale rather than on back-transformed relative expression values. Normality of residuals was assessed and no significant departure was detected. Fold changes and confidence intervals are reported in **Supplementary Table S8.** Summary values are reported as geometric means. Analyses were performed in GraphPad Prism.

## Results

### T6SS1 and T6SS2 make distinct contributions to antibacterial and antifungal competition in Vp_AHPND_ isolates D4 and 297B

To begin our investigation, we identified the *V. parahaemolyticus* strains D4 (Mexico) and 297B (Vietnam) as suitable, geographically distinct representatives of Vp_AHPND_. These isolates have been used in several works, including foundational investigations of AHPND by Nunan et al. (2014) and Han et al. (2015) [10, 11]. Moreover, both D4 and 297B harbor complete T6SS1 and T6SS2 gene clusters, and exhibit antibacterial activity under warm, marine-like conditions (i.e. 30°C, 3% [wt/vol] NaCl) [10–12, 23, 59, 66]. Notably, while the *V. parahaemolyticus* T6SS2 is most active at a lower temperature and salinity range (i.e. 23°C, 1% [wt/vol] NaCl), some strains also employ T6SS2 under the warm, marine-like conditions typically associated with T6SS1 [44].

We first sought to evaluate how inactivation of T6SS1, T6SS2, or both systems would influence interbacterial killing by D4 and 297B. To this end, deletions in the genes encoding Hcp1 and/or Hcp2 were made by allelic exchange, thus rendering one or both systems inoperable [48, 49]. Importantly, retention of the *pirA/B^vp^*locus in mutant derivatives of D4 and 297B was confirmed using the duplex PCR screen described by Han et al. (2015) **(Supplementary Fig. S1a** and **S1b)** [11]. Growth experiments were also performed to ensure that the above mutations did not incur a growth defect **(Supplementary Fig. S2a** and **S2b)**. Mutants of D4 and 297B were then used as attackers in T6SS competition experiments against *E. coli* DH5α prey to assess killing ability relative to the parent strains (**Fig. 2a** and **2b**).

**Fig. 2.**
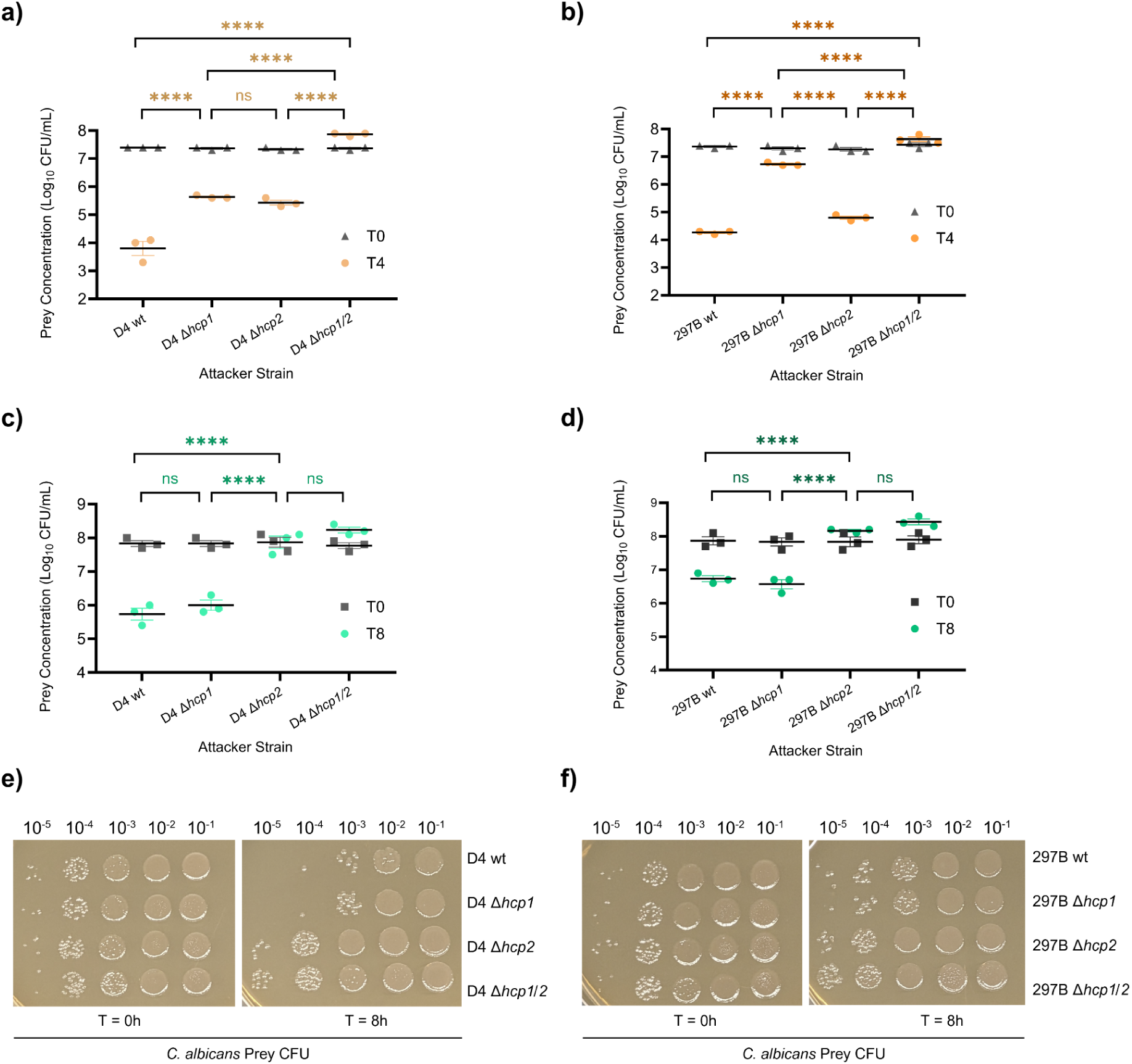
T6SS1 and T6SS2 differentially mediate antibacterial and antifungal competition in Vp_AHPND_ isolates D4 and 297B. (a, **b)** Recovery of *E. coli* DH5α prey before (T0, triangles) and after (T4, orange circles) 4 hour co-incubation with the indicated *V. parahaemolyticus* attacker strain on Marine Luria Bertani at 30°C. Bars show mean ± SEM. **(c**, **d)** Recovery of *C. albicans* ATCC 90028 prey before (T0, squares) and after (T8, green circles) 8 hour co-incubation with the indicated *V. parahaemolyticus* attacker strain on TSA+ at 30°C. The statistical significance between samples at T4 **(a**, **b)** and T8 **(c**, **d)** was determined by two-way ANOVA (matched by biological replicate) with Tukey’s test for multiple comparisons; \*\*\*\**P <* 0.0001, ns = not significant. Values at T0 are not statistically different. **(e**, **f)** Recovery of *C. albicans* 90028 prey before and after co-incubation with Vp_AHPND_ attackers. Serial 10-fold dilutions of prey cells mixed with the indicated attacker strain were spotted onto chloramphenicol-supplemented YPD agar plates at the start (T0) and end (T8) of competition experiments. Plates were incubated for 24 hours at 30°C and then visualized. Symbols in **a**, **b**, **c**, and **d** denote values from individual biological replicates (*n* = 3), each the mean of triplicate platings. Bars indicate mean ± SEM.

Wild type D4 and 297B demonstrated efficient killing of *E. coli* prey under warm, marine-like conditions on a solid surface (i.e., agar media containing 3% [wt/vol] NaCl, with a 4 hour incubation at 30°C) (**Fig. 2a** and **2b**). Deletion of either *hcp1* or *hcp2* alone resulted in attenuated killing ability relative to the parent strains (*P* < 0.0001), whereas double mutants lacking both genes (i.e. D4 Δ*hcp1/2* and 297B Δ*hcp1/2*) failed to decrease *E. coli* prey CFU/mL concentrations after 4 hours. Interestingly, the contributions of T6SS1 and T6SS2 to the antibacterial activity of D4 were comparable under the tested conditions, as indicated by a statistically insignificant (*P* = 0.492) difference in log_10_ CFU/mL of *E. coli* recovered from competitions against D4 Δ*hcp1* and D4 Δ*hcp2*. On the other hand, 297B appears to rely more heavily on T6SS1 for antibacterial killing based on the same metric (*P* < 0.0001).

To ascertain whether the T6SSs of D4 and 297B also enable competition against fungal species, we performed 8 hour contact-killing experiments at 30°C on tryptic soy agar supplemented with 2% [wt/vol] NaCl (TSA+) using *Candida albicans* ATCC 90028 as a prey strain **(Fig. 2c**, **2d**, **2e**, and **2f)** [53]. Remarkably, D4 wild type and D4 Δ*hcp1*, but not D4 Δ*hcp2* or D4 Δ*hcp1/2*, demonstrated the ability to antagonize *C. albicans* (*P* < 0.0001), as marked by average log_10_ declines in prey CFU/mL concentration of 2.1 (D4 wt) and 1.8 (D4 Δ*hcp1*) from T0 to T8. Likewise, co-incubation with either 297B wild type or 297B Δ*hcp1*, but not 297B Δ*hcp2* or 297B Δ*hcp1/2* (*P* < 0.0001), reduced viable *C. albicans* recovery.

Killing of *C. albicans* prey by 297B was less pronounced compared with D4, as reflected in log_10_ CFU/mL declines of only 1.1 (297B wt) and 1.3 (297B Δ*hcp1*). Taken together, these results indicate that both of the tested Vp_AHPND_ isolates antagonize fungal competitors under warm, high salinity conditions in a T6SS2-dependent fashion.

Given the strain-specific differences observed in antibacterial and antifungal activity, we performed a comparative analysis of the T6SS effector/immunity (E/I) repertoires employed by D4 and 297B, beginning with a search for homologs of experimentally confirmed effectors reported in the literature. Inspection of the D4 and 297B genomes (GCF_050888945.2 and GCF_002114585.1 respectively) revealed several putative effectors associated with T6SS1 or T6SS2 (**Table 1)**.

**Table 1.** Predicted T6SS effector toxins encoded by Vp_AHPND_ isolates D4 and 297B. Predicted effector proteins associated with T6SS1 and T6SS2 were identified by sequence homology to known effectors and domain analysis carried out using HHPred and Interpro [54, 56, 57]. Gene locus information is provided for assemblies GCF_050888945.2 (D4), and GCF_002114585.1 (297B). **^a^**, predicted by sequence homology; **^b^**, predicted by Interpro; **^c^**, predicted by HHPred. Full alignment results are presented in the **supplementary information** of this manuscript.

| Predicted association | Predicted activity or domain <sup>a, b, c</sup> | *Vibrio parahaemolyticus* 13-306/D4 |  | *Vibrio parahaemolyticus* 12-297/B |  | Literature homolog |
| --- | --- | --- | --- | --- | --- | --- |
|  |  | Protein accession | Gene locus | Protein accession | Gene locus | Effector name |
| T6SS1 | VP1390-like <sup>a</sup> | WP_085609853.1 | ACMX29_RS08515 | WP_085577065.1 | B5C30_RS15275 | VP1390 |
|  | VP1388-like <sup>a</sup> | WP_024699427.1 | ACMX29_RS08525 | WP_029793730.1 | B5C30_RS15265 | VP1388 |
|  | PAAR-like; AHH nuclease <sup>b</sup> | WP_031852689.1 | ACMX29_RS08395 | WP_237771759.1 | B5C30_RS02625 | VP1415 |
|  | PAAR-like; PoNe nuclease <sup>b</sup> | - | - | WP_029857615.1 | B5C30_RS14465 | PoNe |
|  | Channel-forming toxin<sup>c</sup> | WP_025789713.1 | ACMX29_RS24340 | WP_025789713.1 | B5C30_RS04275 | TseVs |
| T6SS2 | Rhs repeats <sup>b</sup> | WP_191786262.1 | ACMX29_RS07915 | WP_236575995.1 | B5C30_RS19400 | T2Rhs-Nuc <sup>BB22OP</sup> |
|  | Lipase class 3 <sup>b</sup> | WP_020840162.1 | ACMX29_RS12530 | WP_031856000.1 | B5C30_RS10045 | T2LipB <sup>BB22OP</sup> |
|  | Unknown <sup>a</sup> | WP_025550820.1 | ACMX29_RS06995 | - | - | T2Unk |
| | $\alpha/\beta$ hydrolase <sup>b</sup> | WP_228767727.1 | ACMX29_RS20475 | - | - | T2Hydro <sup>RIMD</sup> |
| Unknown | Tme-like <sup>a</sup> | WP_025542139.1 | ACMX29_RS13355 | - | - | Tme <sup>T9109</sup> |
|  | AHH nuclease <sup>b</sup> | WP_025520465.1 | ACMX29_RS18445 | - | - | Va16922 |

Among the identified effectors are VP1415 homologs, which contain PAAR-like and AHH nuclease domains and are encoded a short distance downstream of *dotU* at one end of the T6SS1 gene cluster [23, 32]. Located near the opposite end of the T6SS1 cluster is a tricistronic operon coding for VP1388-1390-like effector modules [43]. In D4, this operon is followed by a gene coding for a large tandem repeat protein (<u>WP_318301207.1</u>) homologous to the putative lipoprotein TasL (<u>WP_135354012.1</u>) from the bobtail squid symbiont *Aliivibrio fischeri* [67]. While not a T6SS cargo molecule in itself, TasL is employed by *A. fischeri* to aggregate competitor cells within the host environment, thereby facilitating the contact requisite for T6SS-dependent killing [67]. As previously reported, 12-297/B encodes the VgrG1b module, which includes the Polymorphic Nuclease effector (PoNe) and immunity protein (PoNi) [66].

Common between D4 and 297B are homologs of the nuclease effector T2Rhs-Nuc (both with PoNe_DUF637 family C-terminal toxin domains), and the lipase T2LipB, which together serve as the “core” substrates delivered by T6SS2 [44]. The predicted T6SS2 arsenal of D4 is further diversified by homologs of the T2Unk and T2Hydro effectors, the latter of which is proposed to function as an α/β hydrolase. Additional E/I modules associated with either T6SS1 or T6SS2 are predicted within the D4 genome, including an orphan Tme-DUF1240 pair and an auxiliary AHH-Imm11 cassette [42, 44].

### The AHPND-associated pVA1-type plasmid harbors an antibacterial effector/immunity pair

Unexpectedly, our survey of the effector landscape in D4 revealed two hypothetical proteins (<u>WP_025789713.1</u> and <u>WP_025789712.1</u>), encoded by the pVA1-type plasmid as a candidate E/I pair. A 4-bp overlap in the corresponding CDS regions (locus tags <u>ACMX29_RS24340</u> and <u>ACMX29_RS24335</u>) indicated that they are likely co-expressed, as is typical for antibacterial E/I pairs. Intriguingly, this genomic region is proximal to the composite transposon which contains the *pirA^vp^/pirB^vp^* genes (**Fig. 3a**) [13–15]. Gene pairs coding for identical proteins are likewise arranged on the pVA1-type plasmids of geographically diverse Vp_AHPND_ isolates such as L2181 (Ecuador), PSU5579 (Thailand), and CGVP22 (Korea), and also occur on similar plasmids from AHPND-causing *V. harveyi* and *V. owensii* strains [14, 16, 68–70]. Although sometimes annotated as a “small-subunit ribosomal protein”, closer inspection of WP_025789713.1 instead supports assignment as a toxin candidate. Structure predictions performed using HHPred suggest that WP_025789713.1 harbors a C-terminal channel-forming domain similar to those found in group A colicins, while PSORTb (version 3.02) predicts localization of WP_025789713.1 to the cytoplasmic membrane (localization score 9.46) [56, 71].

**Fig. 3.**
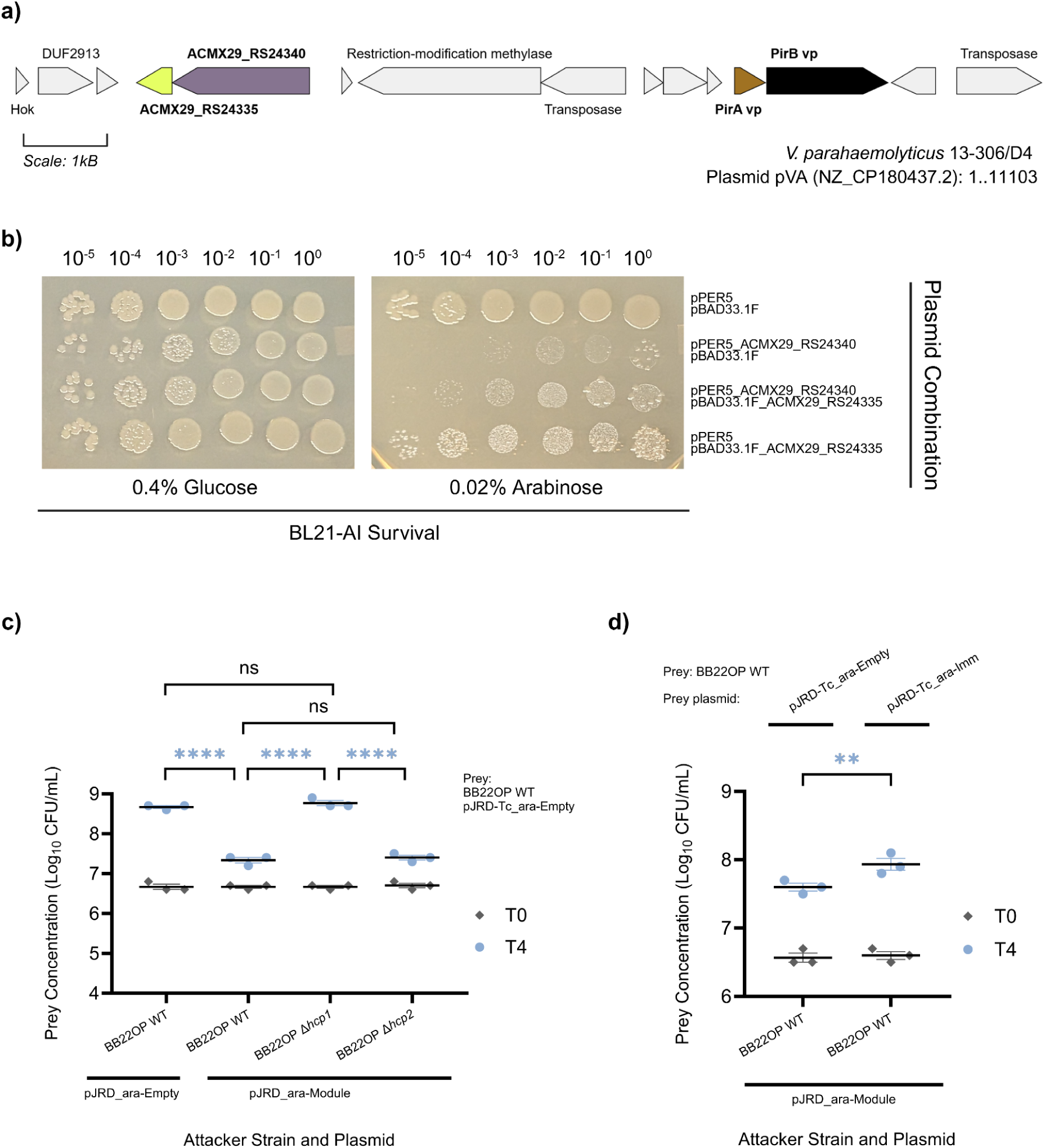
Analysis of WP_025789713.1 and WP_025789712.1 as an E/I pair. **(a)** Genomic context of ACMX29_RS24340 (lead gray) and ACMX29_RS24335 (yellow) on the pVA1-type plasmid of D4 (NZ_CP180437.2). Located nearby is the composite transposon which harbors the CDS for the PirA^vp^ (brown) and PirB^vp^ (black) binary toxin. Image made with Gene Graphics [75]. **(b)** Toxicity from expression of WP_025789713.1 alone or co-expressed with WP_025789712.1 in *E. coli* BL21-AI. Transformants harboring combinations of pPER5, pBAD33.1F, or the indicated expression plasmids were inoculated into LB supplemented with chloramphenicol, kanamycin (for plasmid maintenance), and 0.4% [wt/vol] glucose. Cultures were incubated overnight at 37°C with constant shaking. The next day, cells were pelleted, washed twice with PBS, and normalized to an OD_600_ of 1.0. Serial 10-fold dilutions were spotted onto LB agar plates (pH 8) containing chloramphenicol, kanamycin, and either 0.4% [wt/vol] glucose (to maintain repression) or 0.02% [wt/vol] arabinose (to induce expression). Plates were incubated at 37°C for 24 hours and then inspected to assess growth. **(c**, **d)** Recovery of parental BB22OP prey harboring a tetracycline resistance plasmid (pJRD-Tc_ara-Empty) before (T0, diamonds) and after (T4, blue circles) 4 hour co-incubation with the indicated BB22OP attacker strain on MLB agar supplemented with 0.1% arabinose at 30°C. Attackers carried either an empty MCS control (pJRD_ara-Empty) plasmid or a plasmid expressing the WP_025789713.1/WP_025789712.1 module (pJRD_ara-Module) as indicated below the x-axis. In **(d)**, parental prey containing either pJRD-Tc_ara-Empty or a derivative expressing WP_025789712.1 (pJRD-Tc_ara-Imm) were co-incubated with the indicated attacker on MLB agar supplemented with 0.02% arabinose. Statistical significance between samples at T4 was determined by two-way ANOVA (matched by biological replicate) with Tukey’s test for multiple comparisons **(c)** or Fisher’s LSD test **(d)**; \*\**P* < 0.01, \*\*\*\**P* < 0.0001, ns = not significant. Values at T0 are not statistically different. Symbols in **c** and **d** denote values from individual biological replicates (*n* = 3), each the mean of triplicate platings. Bars indicate mean ± SEM.

A search for similar proteins in the literature revealed homology between WP_025789713.1 and other predicted T6SS effectors (**Table S6**), including the *Pantoea* protein <u>ADU69816.1</u> and the RIX motif protein <u>WP_130243383.1</u> [72, 73]. During the drafting of this manuscript, a report was published describing the *V. cholerae* E1 effector “TseVs” (<u>XDT02158.1</u>), which exhibits robust homology to WP_025789713.1 [74]. Liu et al. (2025) demonstrated ion channel formation and resultant membrane depolarization as the mechanistic basis for TseVs-mediated intoxication, and further showed that TseVs associates with the heterotrimeric VgrG complex of the *V. cholerae* T6SS [74]. Interestingly, Liu et al. (2025) also revealed that the TseVs/TsiVs module imparts a fitness advantage in co-colonization experiments using shrimp shell sections.

Comparison of WP_025789713.1 to TseVs and WP_130243383.1 revealed that the N-terminal region (residues 1-96) of WP_025789713.1 lacks homology to either protein, and does not appear to contain any previously reported loading or adaptor domains. Using the first 96 residues of WP_025789713.1 as a query, we performed four iterations of Position-Specific-Iterated BLAST (PSI-Blast) against the NCBI non-redundant protein database, yielding 63 matches restricted to *Vibrionaceae*. While many of the returned sequences lack domain annotations, the remainder feature putative C-term toxin domains. For example, the *A. fischeri* proteins <u>WP_012534754.1</u> and <u>WP_155656643.1</u> contain Tox-PLDMTX (dermonecrotoxin) and CIF (cycle inhibiting factor) domains respectively, while matches found in *V. owensii* (<u>WP_408738956.1</u>), and *V. coralliilyticus* (<u>WP_234425281.1</u>) harbor nuclease domains associated with polymorphic toxin systems.

To assess whether WP_025789713.1 functions as an antibacterial toxin when delivered to the periplasm, the CDS for ACMX29_RS24340 was cloned into pPER5 (a pBAD derivative), in-frame with a pelB leader sequence, to enable arabinose-inducible expression in *E. coli* BL21-AI [42, 50]. When plated on LB agar (pH 8) supplemented with arabinose, BL21-AI cells transformed with a WP_025789713.1 expression plasmid exhibited steep declines in viability as compared to those harboring an empty pPER5 control (**Fig. 3b**). To evaluate WP_025789712.1 as an antidote to WP_025789713.1-mediated toxicity, WP_025789713.1 and WP_025789712.1 were co-expressed from pPER5 and pBAD33.1F respectively, resulting in a partial rescue of survival. Notably, high-level expression of immunity proteins for periplasmic effectors can prove toxic by itself [42]. For this reason, a low concentration (0.02% [wt/vol]) of arabinose was used in order to mitigate growth defects caused by overexpression of WP_025789712.1.

Our next pursuit was to determine if WP_025789713.1 and WP_025789712.1 constitute an E/I pair associated with either T6SS1 or T6SS2. Antibacterial E/I modules are often confirmed by deleting the corresponding gene pair in a strain of interest, and then using the resulting mutant as prey in competitions against its parental strain [32, 76, 77]. Loss of the immunity gene may render the mutant vulnerable to intoxication when the cognate effector is delivered by parental attackers. Alternatively, predicted E/I gene pairs can be supplied on a plasmid for ectopic expression in an appropriate surrogate attacker, which is then competed against its parental strain [42, 78, 79]. Since ACMX29_RS24340-35 is borne on a pVA1-type plasmid, which is multicopy and maintained by a PSK system, we chose the latter approach [12, 16].

We began by inserting the combined CDS for ACMX29_RS24340-35 into the MCS of pBAD33.1F, in-frame with a C-terminal FLAG tag. The resulting construct was then used as a template for amplification of the ACMX29_RS24340-35 CDS and surrounding pBAD elements (*araC*, the *araBAD* promoter, and *rrnB* terminators) as a single fragment [79]. This fragment was subsequently introduced into the broad host range plasmid pJRD215 to enable arabinose-inducible expression of WP_025789713.1/WP_025789712.1 in the non-AHPND *V. parahaemolyticus* strain BB22OP, which otherwise lacks this module [51, 80].

Wild type and *hcp* mutant strains of BB22OP armed with either the WP_025789713.1/WP_025789712.1 expression plasmid (pJRD_ara-Module) or an empty MCS control (pJRD_ara-Empty) were then competed against parental BB22OP harboring a tetracycline resistance plasmid (pJRD-Tc_ara-Empty; for enumeration on tetracycline containing MMM plates). Attacker and prey cells were mixed in a 10:1 ratio (attacker-to-prey) and spotted onto MLB agar plates containing 0.1% arabinose to induce expression from the *araBAD* promoter. As before, competition plates were incubated at 30°C, and prey CFU before (T0) and after (T4) coincubation were determined by plating serial 10-fold dilutions on selective media (**Fig. 3c**).

In contrast to attackers harboring an empty MCS control plasmid, wild type attackers equipped with the pJRD_ara-Module plasmid gained the ability to inhibit growth of parental prey, as indicated by differences in Log_10_ CFU/mL concentrations of parental BB22OP at T4 (*P* < 0.0001). Interestingly, deletion of *hcp1*, but not *hcp2*, rendered attackers with the pJRD_ara-Module plasmid unable to antagonize parental prey, with no significant difference (*P* = 0.5705) in concentrations of prey recovered from competitions against wild type attackers with a control plasmid, or Δ*hcp1* attackers with pJRD_ara-Module.

To assess protection conferred by the suspected immunity protein WP_025789712.1, parental prey were equipped with a plasmid to enable arabinose-inducible expression of the ACMX29_RS24335 CDS (pJRD-Tc_ara-Imm), and competed against wild type attackers armed with pJRD_ara-Module on MLB agar plates supplemented with 0.02% arabinose (**Fig. 3d**). Compared to prey containing an empty MCS control (pJRD-Tc_ara-Empty), prey that were given the WP_025789712.1 expression plasmid exhibited significantly higher (*P* = 0.0090) Log_10_ CFU/mL concentrations after coincubation with BB22OP wild type attackers expressing the WP_025789713.1/WP_025789712.1 module. Taken together, these results indicate that WP_025789713.1 and WP_025789712.1 comprise a functional E/I pair which, in *V. parahaemolyticus*, facilitate competitive interactions in a T6SS1-dependent manner.

### T6S is necessary for full virulence of Vp_AHPND_ in *L. vannamei* postlarvae under warm, high-salinity conditions

To examine whether T6SS1 and T6SS2 contribute to virulence during shrimp infection by our Vp_AHPND_ strains, we employed an immersion bioassay adapted from Tran et al. (2013), using the commercial shrimp species *L. vannamei* as a model [9]. Disease-free *L. vannamei* were obtained from Miami Aquaculture (Boynton Beach, Florida, USA) and maintained in a filtered and aerated aquarium tank containing Artificial Seawater (ASW). For infection challenges, *L. vannamei* postlarvae (PL) ranging from PL24–PL32 were distributed according to treatment group and immersed in either sterile ASW (inoculated with PBS) or ASW suspensions containing approximately 1×10^8^ CFU/mL of the indicated bacterial strain. After a 15 minute incubation at 30°C, PL were transferred to experiment containers filled with sterile ASW (30 ppt salinity) and inoculated with either a PBS control, or a sufficient volume of bacterial suspension to yield a concentration of approximately 1×10^6^ CFU/mL. Experiment containers were maintained at a temperature of 30°C with aeration provided by sterile airstones, and PL survival was determined at 12, 24, and 48 hours post-infection (**Fig. 4a**, **4b**, and **4c)**.

**Fig. 4.**
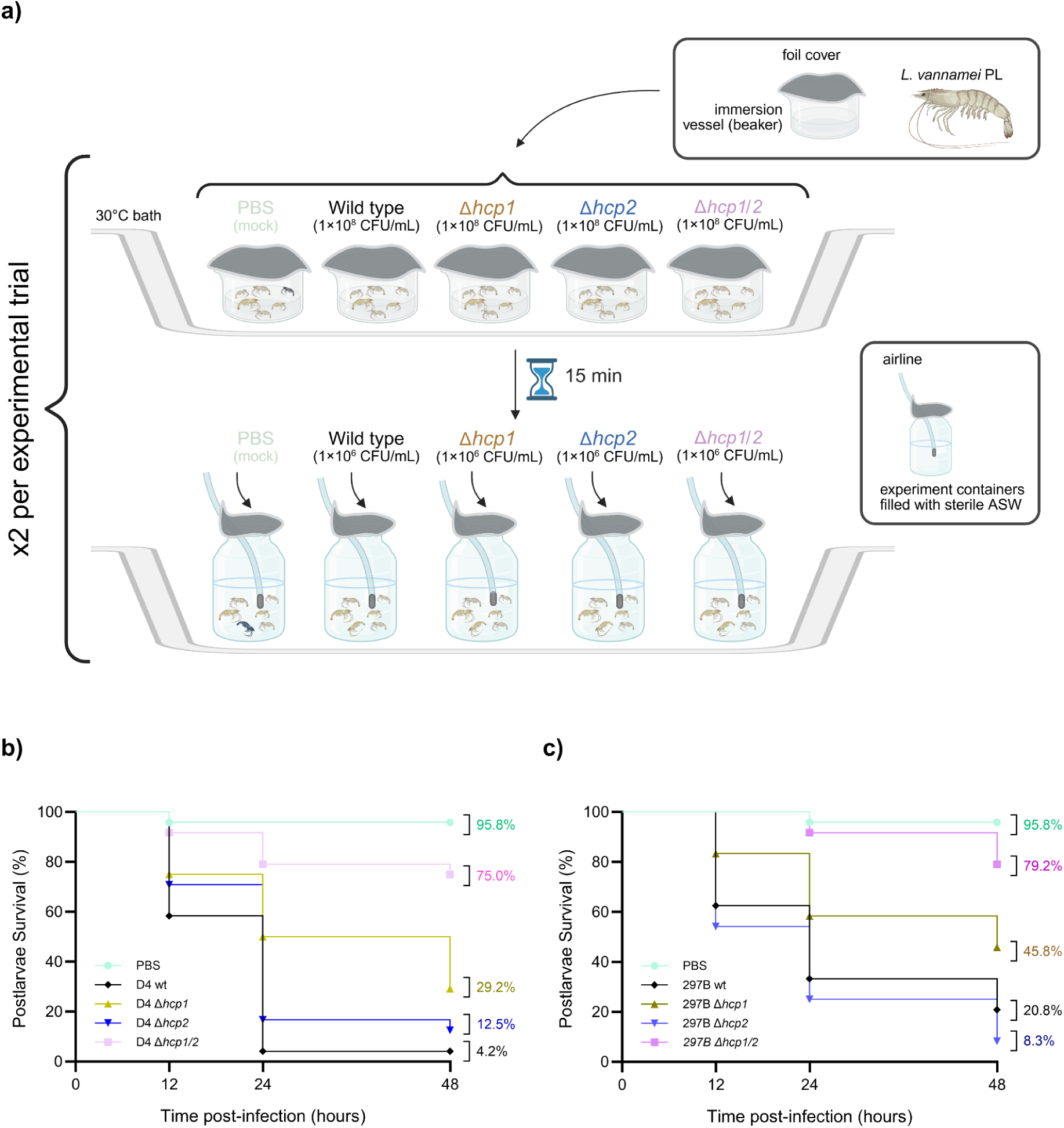
Type VI secretion is required for full virulence of D4 and 297B in *L. vannamei* postlarvae. **(a)** Schematic of the infection bioassay used to assess Vp_AHPND_ virulence in *L. vannamei*. Postlarval *L. vannamei* (PL24–PL32) were allocated into beakers containing sterile ASW, and inoculated with sterile PBS or a sufficient volume of the indicated *V. parahaemolyticus* strain to achieve a concentration of approximately 1×10^8^ CFU/mL. After a 15 minute immersion at 30°C, PL were transferred into foil-covered experiment containers filled with 400 mL of sterile ASW (30°C; 30 ppt salinity) and reinoculated with either PBS or a volume of bacterial suspension to yield a final density of approximately 1×10^6^ CFU/mL. Shrimp were fed 10% of their estimated body weight once daily, and survival rates were assessed at 12, 24, and 48 hours post-infection. **(b**, **c)** Kaplan-Meier survival curves for postlarvae infected with wild type or T6SS-deficient derivatives of D4 **(b)** or 297B **(c)** under warm, high-salinity conditions (30°C, 30 ppt salinity). Cumulative survival at 48 h is indicated beside each curve. Statistical differences between groups were determined by log-rank test. Data shown represent two independent experiments, each performed using 2 replicate containers for every treatment group, with 6 shrimp per container (*N* = 24 total animals per treatment group).

Shrimp infected with wild type D4 or 297B exhibited significantly reduced survival compared to those inoculated with a PBS control (*P* < 0.0001), with cumulative mortalities approaching 100% (D4 wt) or 80% (297B wt) within 48 hours. Neither D4 Δ*hcp2* (*P* = 0.203) nor 297B Δ*hcp2* (*P* = 0.295) were significantly attenuated in causing *L. vannamei* mortality compared with the respective parent strains.

Contrastingly, inactivation of T6SS1 alone impaired killing of *L. vannamei* by D4 Δ*hcp1* (*P* = 0.0045) and 297B Δ*hcp1* (*P* = 0.0437), and resulted in higher cumulative PL survival at 48 hours post-infection.

Moreover, simultaneous disruption of both T6SS1 and T6SS2 further compromised the virulence of both isolates, as marked by significant differences in *L. vannamei* survival when challenged with either D4 Δ*hcp1/2*, 297B Δ*hcp1/2*, or the corresponding parent strains (*P* < 0.0001). Notably, *L. vannamei* inoculated with these mutants exhibited average survival rates of approximately 75% (D4 Δ*hcp1/2*) and 79% (297B Δ*hcp1/2*) at 48 hours post-infection. Dead and moribund PL challenged with D4, 297B, or mutant derivatives displayed gross morphological and behavioural signs classically associated with AHPND, namely discoloration and atrophy of the hepatopancreas [81]. Individuals that were putatively infected but still alive displayed lethargy punctuated by weak, “corkscrew” swimming, as others have described [82].

### Antibiotic-mediated reduction of host bacteria partially compensates the virulence defect of D4 T6SS mutants

Various works have proposed that T6SS1 is involved in competition against host-associated microbes during shrimp infection [23, 24, 26, 27]. Prompted by this, we conducted additional challenge experiments with D4 against *L. vannamei* PL harboring lower titers of culturable bacteria (**Fig. 5a** and **5b**). Due to the lack of an axenic or gnotobiotic *L. vannamei* model, antibiotic-treated *L. vannamei* have previously been employed as an alternative [83]. Taking a similar approach, we began by identifying an antibiotic cocktail that would be suitable for reducing the *L. vannamei* midgut bacteriome without adversely impacting the survival of uninfected PL. Leveraging the innate drug resistance profile of D4, a formula consisting of tetracycline (10 µg/mL), colistin (10 µg/mL), ampicillin (100 µg/mL), and vancomycin (50 µg/mL) was chosen [59–61].

**Fig. 5.**
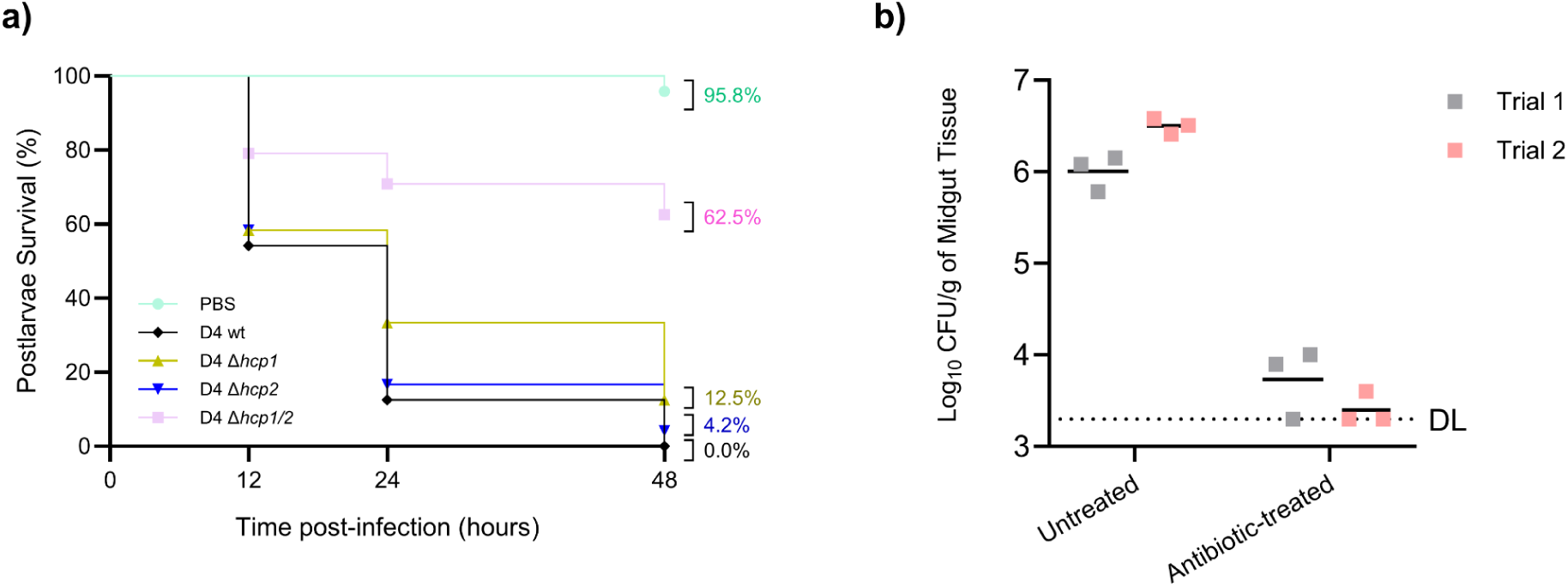
Pre-treatment of *L. vannamei* postlarvae with an antibiotic cocktail reduces culturable host-associated bacteria and partly masks the attenuated virulence of D4 Δ*hcp1*. **(a)** Kaplan-Meier survival curves for antibiotic-treated *L. vannamei* postlarvae infected with wild type D4 or the indicated T6SS mutants. Cumulative survival at 48 h is indicated beside each curve. Statistical differences between groups were determined by log-rank test. Data shown represent two independent experiments, each performed using 2 replicate containers for every treatment group, with 6 shrimp per container (*N* = 24 total animals per treatment group). **(b)** Recovery of culturable bacteria from midgut tissue samples of untreated and antibiotic-treated postlarvae, expressed as Log_10_ CFU/g. Results from independent experimental trials are shown in gray (Trial 1) and pink (Trial 2); each square represents one midgut sample. The dotted line indicates the limit of detection (DL). Values below the limit of detection were assigned the DL for plotting and summary calculations [84].

To measure the impact of antibiotic administration with respect to culturable bacteria present in the *L. vannamei* midgut, hepatopancreases and intestines were aseptically harvested from antibiotic-treated and -untreated shrimp, weighed, and then homogenized into sterile ASW. Estimated bacterial titers were obtained by spotting serial 10-fold dilutions of the resulting homogenates onto MLB agar plates, which were then incubated for 24 hours (30°C) prior to inspection. For infection experiments using antibiotic-treated shrimp, PL were distributed into experiment beakers containing ASW (30 ppt salinity) supplemented with the aforementioned antibiotic cocktail and incubated for 24 hours at 30°C. Afterward, antibiotic-treated ASW was exchanged with sterile, untreated ASW to remove leftover drug residues, and immersion bioassays were carried out as before.

Incubation in ASW supplemented with the antibiotic mixture reduced average culturable bacterial loads recovered from postlarval midgut tissues by approximately 2.3 and 3.1 log_10_ CFU/g in two independent trials (**Fig. 5b**). Moreover, in challenges of antibiotic-treated *L. vannamei*, post-infection mortalities caused by D4 Δ*hcp1* were not found to be significantly different (*P* = 0.1123) from those caused by the wild type strain (**Fig. 5a**). On the other hand, statistical differences in survival between shrimp infected with D4 Δ*hcp1/2* and those infected with D4 wild type were observed (*P* < 0.0001). When interpreting these data, it is important to note limitations imposed by our choice of model compared to a true axenic or gnotobiotic system. Namely, CFU determinations made by spot-plating reveal only a fraction of viable bacteria present in the shrimp midgut, and do not account for viable but nonculturable (VBNC) organisms or species which are unable to form colonies under the tested media and incubation conditions [85–87].

### Inactivation of T6S does not suppress *pirA/B^vp^* transcription in D4

Past studies of T6S have revealed that deletion of T6SS structural genes can incur various unanticipated phenotypic changes and influence the expression of many non-T6SS genes [38, 88–91]. We therefore chose to investigate whether T6SS inactivation, as achieved through deletion of *hcp1* and/or *hcp2*, impacts expression of the *pir* genes in D4. Using primer pairs developed by Lin et al. (2022), we performed RT-qPCR to measure transcription levels of *pirA^vp^* and *pirB^vp^*in log-phase cultures of D4 wild type and mutant strains grown in TSB+ at 30°C (**Fig. 6**) [65].

**Fig. 6.**
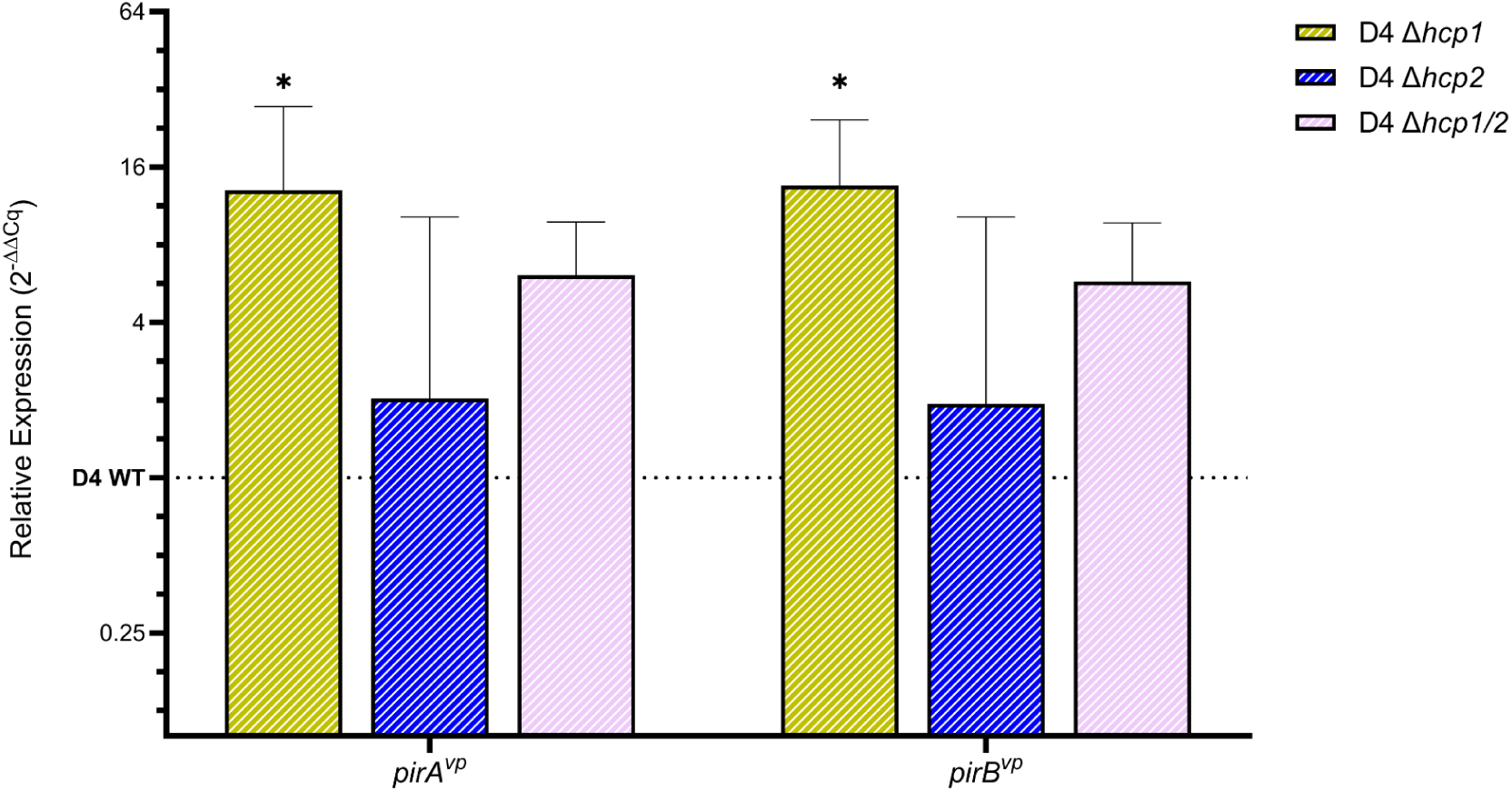
D**e**letion **of the *hcp* genes does not hamper transcription of *pirA/B^vp^* in D4 under warm, high salinity conditions.** RT-qPCR analysis of *pirA^vp^* and *pirB^vp^* expression in wild type and *hcp* mutant strains of D4 during log-phase growth in TSB+ at 30 °C, using *gyrB* as the reference gene. Bars show relative expression (2^−ΔΔCq^) in each mutant, with reference to wild type D4 (indicated by a dashed line at 1), on a log_2_ axis. Bars representing the geometric mean of three biological replicates (*n* = 3) are shown. Error bars span the geometric mean multiplied and divided by the geometric SD factor. Statistical significance was evaluated independently for *pirA^vp^*and *pirB^vp^* using separate one-way ANOVAs on ΔC_q_ values followed by Dunnett’s multiple comparisons against D4 wild type; \**P* < 0.05.

Global one-way ANOVA of *pirA^vp^* transcript levels did not demonstrate a significant effect of genotype on expression (*F*(3, 8) = 3.473, *P* = 0.0707). However, Dunnett’s multiple comparisons test identified increased *pirA^vp^* transcript abundance in D4 Δ*hcp1* relative to wild type D4 (*P* = 0.0436). The mean ΔC_q_ for *pirA^vp^* decreased from −6.33 in wild type D4 compared to −10.05 in D4 Δ*hcp1* (ΔΔC_q_ = −3.72). Since ΔC_q_ is calculated as C_q_(*pirA^vp^*) − C_q_(*gyrB*), this decrease corresponds to an approximately 13-fold increase in relative expression, although there is a wide confidence interval with a lower bound near one (95% CI 1.1–160). The D4 Δ*hcp2* (2.0-fold, 95% CI 0.17–25, *P* = 0.7574) and D4 Δ*hcp1/2* (6.1-fold, 95% CI 0.50–74, *P* = 0.1642) mutants showed point estimates in the same direction but they did not differ significantly from the wild type. Genotype accounted for approximately 56.6% of the variance in *pirA^vp^*expression (*R*^2^ = 0.5657).

The variation in *pirB^vp^* transcript levels across the four strains also did not reach statistical significance by global ANOVA (*F*(3, 8) = 3.678, *P* = 0.0625), although the Dunnett’s test identified increased *pirB^vp^* transcript abundance in D4 Δ*hcp1* relative to wild type D4 (14-fold, 95% CI 1.2–157, *P* = 0.0383). The D4 Δ*hcp2* (2.0-fold, 95% CI 0.17–23, *P* = 0.7740) and D4 Δ*hcp1/2* (5.8-fold, 95% CI 0.50–68, *P* = 0.1642) mutants showed point estimates in the same direction but were not significantly different than wild type. Genotype accounted for approximately 58.0% of the variance in *pirB^vp^* expression (*R*^2^ = 0.5797). The estimated fold changes for *pirA^vp^* and *pirB^vp^*were comparable in every mutant background, and no comparison in either gene indicated reduced transcript abundance relative to wild type.

## Discussion

Contact-killing experiments indicate that D4 and 297B employ both T6SS1 and T6SS2 to varying degrees against bacterial competitors under warm, high salinity conditions. In 297B, antibacterial activity appears to be more strongly correlated with T6SS1 function, which may reflect differences in effector arsenal between this strain and D4. Surprisingly, both isolates also demonstrate the ability to kill fungal prey in a T6SS2-dependent manner, and may therefore utilize this system to compete against fungi present in the shrimp gut and surrounding waters [92, 93]. Genomic interrogation of D4 and 297B indicates that these strains harbor overlapping yet distinct T6SS effector arsenals, with notable differences including a VgrG1b module present in 297B, and a greater abundance of T6SS2 effectors in D4 [23, 44, 66]. However, it is unclear whether the antifungal phenotypes shown by D4 and 297B are mediated by homologs of known T6SS2 effectors with potential broad-spectrum activity (e.g. nucleases).

Given the many challenges which impede T6SS effector studies, it can be reasoned that D4 and 297B may host additional effectors that await discovery and characterization. Highlighting this, our in silico investigations revealed a previously unrecognized E/I cassette neighboring the *pirAB*-Tn903 locus on pVA1-type plasmids. Functional experiments show that upon acquiring and expressing this locus, a non-AHPND surrogate attacker (*V. parahaemolyticus* BB22OP) gains the ability to antagonize its parental strain in a T6SS1-dependent manner. Intriguingly, Liu et al. (2025) recently demonstrated that a homologous E/I pair found primarily in non-toxigenic strains of *V. cholerae* grants a competitive edge over toxigenic rivals during co-colonization of shrimp shells [74]. We hypothesize that this ecological dynamic could be reversed in the case of *V. parahaemolyticus* strains, which may derive a similar advantage from acquiring pVA1-type plasmids that also harbor the *pir* toxin genes.

Mortality data from the *L. vannamei* challenge experiments indicate that perturbation of T6SS1 modestly attenuates the virulence of D4 and 297B against *L. vannamei* PL in a warm, high-salinity environment. Opposingly, within the context of a functional T6SS1, inactivation of T6SS2 did not significantly reduce mortalities caused by D4 and 297B under the same conditions. However, simultaneous disruption of both T6SSs resulted in steeply impaired virulence in both isolates, which may suggest partial overlap in the contributions of T6SS1 and T6SS2 towards pathogenicity. Survival comparisons were made at the level of individual shrimp, and we thus acknowledge potential container-level effects as a limitation of our statistical approach. Importantly, we chose water temperature and salinity conditions that are applicable to *L. vannamei* aquaculture systems and also conducive to T6SS activity in *V. parahaemolyticus* D4 and 297B [23, 39, 94–96]. To simulate the typical occurrence of AHPND outbreaks within 20 to 35 days after stocking of shrimp ponds, *L. vannamei* ranging from PL24 to PL32 were used for infections [1–3]. We propose that modifying experiment parameters such as water temperature or salinity may influence the relative contributions of T6SS1 and T6SS2 to Vp_AHPND_ pathogenicity. For instance, T6SS2 may play a greater role in virulence under lower temperature and salinity conditions (e.g., 23–26°C and 10–20 ppt salinity).

Hypotheses concerning the role of T6S during AHPND progression have focused on the ability of Vp_AHPND_ to kill bacterial competitors within the host [23, 24, 26, 27, 45]. As a preliminary evaluation of this model, we conducted challenge experiments using *L. vannamei* that had been pre-treated with an antibiotic cocktail to reduce culturable bacterial loads occurring in the midgut [83]. In antibiotic-treated *L. vannamei*, cumulative mortality rates produced by D4 Δ*hcp1* and wild type D4 were not statistically different (*P* = 0.1123), but remained low in challenge experiments with D4 Δ*hcp1/2* as reflected by an average survival rate of >60% at the 48 h timepoint. Notably, conclusions that can be drawn from these results are limited by our substitution of antibiotic-treated shrimp in place of a true axenic or gnotobiotic model. While the present study may support competitive displacement as a possible function of T6SS1 during AHPND pathogenesis, additional metagenomic evidence is needed in order to confirm or dismiss related speculations made in the literature. Further, we are unable to confirm antagonism against host microbes as the sole explanation for T6S-dependent virulence phenotypes exhibited by D4 and 297B.

Despite long being regarded exclusively as toxin delivery platforms, mounting evidence has shown that T6SSs engage in a wider range of processes than once believed [36, 37, 97, 98]. Supporting this notion is the observation that mutations affecting T6SS function can produce many unexpected phenotypic and regulatory consequences [38, 88–91]. To address the possibility that deletion of the *hcp* genes hampers virulence through pleiotropic effects on *pirA/B^vp^* expression, we employed RT-qPCR to measure the abundance of *pirA^vp^* and *pirB^vp^* transcripts in wild type and mutant derivatives of D4 [65]. All comparisons between the D4 wild type strain and *hcp* mutant derivatives, across both target genes, yielded point estimates indicating increased rather than decreased transcript abundance, with D4 Δ*hcp1* showing statistically significant increases for *pirA^vp^* (*P* = 0.0436) and *pirB^vp^*(*P* = 0.0383). Accounting for the number of biological replicates and the width of the associated confidence intervals, we interpret these data as establishing the absence of transcriptional suppression rather than as quantifying the magnitude of any increase.

Here we offer empirical evidence for T6S as a pathogenicity factor during shrimp infection by the Vp_AHPND_ isolates D4 and 297B under warm, high-salinity conditions. Although the mechanisms underpinning T6S-dependent virulence await further clarification, we cautiously support displacement of the *L. vannamei* bacteriome as one of the functions mediated by the *V. parahaemolyticus* T6SS1 during host colonization. Additionally, we identify an antibacterial E/I gene pair disseminated on pVA1-type plasmids which may enhance the competitive fitness of Vp_AHPND_ isolates in a T6SS1-dependent fashion. Given past reports of pleiotropic effects arising from T6SS inactivation, we also chose to investigate possible consequences of *hcp* deletion on *pirA/B^vp^*transcription. Gene expression analysis by RT-qPCR does not qualify a model in which the reduced virulence of D4 Δ*hcp1* and Δ*hcp1/2* is driven by transcriptional suppression of *pirA/B^vp^*. Intriguingly, the T6SSs of *V. parahaemolyticus* have been proposed to serve additional functions apart from contact-dependent antagonism. Studies by Yu et al. (2012) and Pinkerton et al. (2019) suggest that T6SS1 and T6SS2 contribute towards adhesion of *V. parahaemolyticus* to host cells, while more recent works implicate T6SS2 as a determinant of biofilm formation in some strains [25, 99–101]. However, the relevance of these findings to Vp_AHPND_ pathogenesis under warm, high-salinity conditions has not yet been demonstrated. Moreover, a complete inventory of T6SS cargo molecules deployed by *V. parahaemolyticus* remains to be defined. Further progress in this direction may therefore grant new insights concerning the scope of T6SS functionality in Vp_AHPND_.

