## Supplementary Information for "Type VI Secretion Promotes Microbial Antagonism and Contributes to Pathogenesis by AHPND-causing *Vibrio parahaemolyticus*"

**Table S1 Strains used in this study**

| Strain name | Genotype [a] | Notes | Source |
|---|---|---|---|
| *Vibrio parahaemolyticus* 13-306/D4 | Wild type | An AHPND-causing *Vibrio parahaemolyticus* isolate from Mexico. Abbreviated as D4. | Obtained from Kim Orth |
| *Vibrio parahaemolyticus* 13-306/D4 Δ*hcp1* | ΔACMX29_RS08500 | A D4 derivative with an in-frame deletion of the *hcp1* gene. | This study |
| *Vibrio parahaemolyticus* 13-306/D4 Δ*hcp2* | ΔACMX29_RS24045 | A D4 derivative with an in-frame deletion of the *hcp2* gene. | This study |
| *Vibrio parahaemolyticus* 13-306/D4 Δ*hcp1/2* | ΔACMX29_RS08500 ΔACMX29_RS24045 | A D4 derivative with in-frame deletions of the *hcp1* and *hcp2* genes. | This study |
| *Vibrio parahaemolyticus* 12-297/B | Wild type | An AHPND-causing *Vibrio parahaemolyticus* isolate from Vietnam. Abbreviated as 297B. | Obtained from Kim Orth |
| *Vibrio parahaemolyticus* 12-297/B Δ*hcp1* | ΔB5C30_RS15290 | A 297B derivative with an in-frame deletion of the *hcp1* gene. | This study |
| *Vibrio parahaemolyticus* 12-297/B Δ*hcp2* | ΔB5C30_RS16745 | A 297B derivative with an in-frame deletion of the *hcp2* gene. | This study |
| *Vibrio parahaemolyticus* 12-297/B Δ*hcp1/2* | ΔB5C30_RS15290 ΔB5C30_RS16745 | A 297B derivative with in-frame deletions of the *hcp1* and *hcp2* genes. | This study |
| *Vibrio parahaemolyticus* BB22OP | Wild type | A non-AHPND environmental strain of *Vibrio parahaemolyticus*. | Obtained from Dor Salomon |
| *Vibrio parahaemolyticus* BB22OP Δ*hcp1* | ΔVPBB_RS06665 | A BB22OP derivative with an in-frame deletion of the *hcp1* gene. | Obtained from Dor Salomon |
| *Vibrio parahaemolyticus* BB22OP Δ*hcp2* | ΔVPBB_RS19920 | A BB22OP derivative with an in-frame deletion of the *hcp2* gene. | Obtained from Dor Salomon |
| *Escherichia coli* S17-1 (λ-pir) | Laboratory strain containing λ-pir | Used for maintenance and conjugation of pDM4 and derived plasmids. | Obtained from Kim Orth |
| *Escherichia coli* DH5α | Laboratory strain | Used for transformations and plasmid maintenance. A DH5α strain transformed with an empty pBAD33.1 plasmid was used as a prey strain for T6SS competition assays. | Vendor |
| *Escherichia coli* NEB5α | Commercial strain derived from DH5α | Used for transformations and plasmid maintenance. | Vendor |
| *Escherichia coli* BL21-AI | Commercial strain derived from BL21 | Used for ectopic expression assays. | Vendor |
| *Candida albicans* ATCC 90028 | Laboratory strain | Used as fungal prey in competition assays. | Vendor |

[a] Locus accessions are provided for assemblies GCF_050888945.2 (D4), and GCF_002114585.1 (297B)

**Table S2 Plasmids used in this study**

| Plasmid Name | Description | Use | Source |
|---|---|---|---|
| pDM4 | A CmR, SacB, and oriR6K plasmid which acts as a suicide vector in *Vibrio parahaemolyticus*. | Used to generate in-frame gene deletions via allelic exchange mutagenesis in *Vibrio parahaemolyticus*. | Obtained from Dor Salomon |
| pDM4_Δ*hcp1* (D4) | A pDM4 derivative harboring 1-kb sequences directly upstream and downstream of the *hcp1* gene in D4. | Used to delete *hcp1* in *Vibrio parahaemolyticus* D4. Flanking sequences were cloned into SacI-digested pDM4 via Gibson Assembly. | This study |
| pDM4_Δ*hcp2* (D4) | A pDM4 derivative harboring 1-kb sequences directly upstream and downstream of the *hcp2* gene in D4. | Used to delete *hcp2* in *Vibrio parahaemolyticus* D4. Flanking sequences were cloned into SacI-digested pDM4 via Gibson Assembly. | This study |
| pDM4_Δ*hcp1* (297B) | A pDM4 derivative harboring 1-kb sequences directly upstream and downstream of the *hcp1* gene in 297B. | Used to delete *hcp1* in *Vibrio parahaemolyticus* 297B. Flanking sequences were cloned into SacI-digested pDM4 via Gibson Assembly. | This study |
| pDM4_Δ*hcp2* (297B) | A pDM4 derivative harboring 1-kb sequences directly upstream and downstream of the *hcp2* gene in 297B. | Used to delete *hcp2* in *Vibrio parahaemolyticus* 297B. Flanking sequences were cloned into SacI-digested pDM4 via Gibson Assembly. | This study |
| pBAD33.1 | A plasmid harboring a CmR cassette. | Used to select for *Escherichia coli* DH5α prey on chloramphenicol-containing plates. | Obtained from Dor Salomon |
| pBAD33.1F | A pBAD33.1 derivative with a FLAG tag sequence located downstream of the MCS. | Used for arabinose-inducible expression. | Obtained from Dor Salomon |
| pPER5 | A pBAD derivative with a pelB sequence inserted downstream of the araBAD promoter. Gene products fused to an N-term pelB sequence localize to the periplasm. Contains a KmR cassette. | Used for arabinose-inducible expression of proteins targeted to the periplasm. | Obtained from Dor Salomon |
| pPER5_WP_025789713.1 | A pPER5 derivative containing the CDS for WP_025789713.1 (locus tag ACMX29_RS24340) fused to an N-term pelB and C-term myc-His sequence. | Used for expression and periplasmic localization of WP_025789713.1 in *Escherichia coli* BL21-AI. | This study |
| pBAD33.1F_WP_025789712.1 | A pBAD33.1F derivative containing the CDS for WP_025789712.1 (locus tag ACMX29_RS24335) fused with a C-term FLAG tag sequence. | Used for expression of WP_025789712.1 in *Escherichia coli* BL21-AI. | This study |
| pBAD33.1F_Module | A pBAD33.1F derivative containing the CDS for WP_025789713.1 and WP_025789712.1 (locus tags ACMX29_RS24340-35) fused with a C-term FLAG tag sequence. | Used as an intermediate during construction of pJRD_ara-Module. | This study |
| pJRD215 | A broad host range, RSF1010-family plasmid harboring KmR and SmR resistance markers. | Used as a vector backbone for ectopic expression of genes in *Vibrio parahaemolyticus* BB22OP. | Vendor |
| pCM62 | A broad host range, RK2-family plasmid harboring TetA and TetR. | Used to amplify a TetA-TetR cassette for insertion into pJRD215 digested with XhoI and BmtI. | Vendor |
| pJRD215-Tc | A pJRD215 derivative in which the KmR cassette has been replaced by a TetA-TetR cassette from pCM62. | Used as an intermediate during construction of pJRD-Tc_ara-Empty and pJRD-Tc_ara-Imm. | This study |
| pJRD_ara-Empty | A derivative of pJRD215 containing the pBAD expression machinery and an empty | Used as a control plasmid for *Vibrio parahaemolyticus* BB22OP attackers. | This study |

| | | | |
|---|---|---|---|
| | MCS from pBAD33.1F. | | |
| pJRD_ara-Module | A derivative of pJRD215 containing the pBAD expression machinery and ACMX29_RS24340-35 insert from pBAD33.1F_Module. | Used to equip *Vibrio parahaemolyticus* BB22OP attackers with an arabinose-inducible WP_025789713.1/WP_025789712.1 module. | This study |
| pJRD-Tc_ara-Empty | A derivative of pJRD215-Tc containing the pBAD expression machinery and an empty MCS from pBAD33.1F. | Used as a control plasmid for *Vibrio parahaemolyticus* BB22OP prey to enable selection on tetracycline-containing plates. | This study |
| pJRD-Tc_ara-Imm | A derivative of pJRD215-Tc containing the pBAD expression machinery and ACMX29_RS24335 insert from pBAD33.1F_WP_025789712.1. | Used to equip *Vibrio parahaemolyticus* BB22OP prey with an arabinose-inducible WP_025789712.1 cassette. | This study |

### Table S3 Primers used in this study

| Primer Name | Sequence [a] | Used for | Notes |
|---|---|---|---|
| Vp hcp1 up fwd | tgtggaatcccgggagagctGTCTGTCGTGAACTTGCTCAGTG | Used to amplify 1kb flanking fragments for construction of pDM4_Δ*hcp1* and pDM4_Δ*hcp2*. Upstream forward and downstream reverse primers were used to identify allelic exchange mutants via band size comparison. | Compatible for Gibson Assembly into pDM4 linearized with SacI-HF. Up fwd and down rev primers were also used to screen secondary recombinants for gene deletion. |
| Vp hcp1 up rev | caaaaagcaaCGCTATTTCCTTTTCTAAAATCTGTTTTGTTTAAATCACC | | |
| Vp hcp1 down fwd | ggaaatagcgTTGCTTTTTGCGTAAAGATTCAGGGC | | |
| Vp hcp1 down rev | gcatgcgggtaacctgagctCACTGATGTTAAAGCTGTAACCAGAC | | |
| Vp hcp2 up fwd | tgtggaatcccgggagagctTTTTTATTAACTGGCCCTAGTGGTGTC | | |
| Vp hcp2 up rev | ccgcacaataaaGCTAATCTCCTAGAGCATTATTAATTGACATATTTCG | | |
| Vp hcp2 down fwd | taggagattagcTTTATTGTGCGGAGGGGTTATCTCC | | |
| Vp hcp2 down rev | gcatgcgggtaacctgagctTTTTGAACTTAAAACATTTACCCACAGAAAAGGTTCG | | |
| pirA 284F | TGACTATTCTCACGATTGGACTG | Standard primers for duplex PCR detection of *pirAvp* and *pirBvp* in *Vibrio parahaemolyticus*. Used to confirm the presence of *pirA/Bvp* in mutants of D4 and 297B. | Primers and reaction settings described by Han et al. (2015). |
| pirA 284R | CACGACTAGCGCCATTGTTA | | |
| pirB 392F | TGATGAAGTGATGGGTGCTC | | |
| pirB 392R | TGTAAGCGCCGTTTAACTCA | | |
| pPER5_fwd | aagcttgggcccgaacaaaaactcatc | Used for PCR linearization of pPER5. | Gibson Assembly primers to linearize pPER5 for insertion of CDS in-frame with an N-term pelB and C-term myc-His sequence. |
| pPER5_rev | ggccatcgccggctgg | | |
| pBAD33.1F fwd | gattacaaggatgacgacgataagtgaaag | Used for PCR linearization of pBAD33.1F. | Gibson Assembly primers to linearize pBAD33.1F for insertion of CDS in-frame with a C-term FLAG tag sequence. |
| pBAD33.1F rev | atgtatatctccttcttaaagttaaacaaaattatttctagaggatc | | |
| Tox_pVA Peri fwd | cgctgcccagccggcgatggccATGATCTATAAAACGAAACCATTATTTAACGTTAAGCAAC | Used to generate pPER5_WP_025789713.1. | Compatible for Gibson Assembly into pPER5. |
| Tox_pVA Peri rev | gtttttgttcgggcccaagcttTAACACTTCTTCTTCAATATACTCCGACAGTTC | | |
| 33.1F Tox_pVA fwd | tttaagaaggagatatacatATGATCTATAAAACGAAACCATTATTTAACGT | Used to generate pBAD33.1F_WP_025789712.1 and pBAD33.1F_Module. | Compatible for Gibson Assembly into pBAD33.1F. |
| 33.1F Imm_pVA fwd | ctttaagaaggagatatacatATGAGTTGGTATGAGATGATTGCCAT | | |
| 33.1F Imm_pVA rev | tcgtcgtcatccttgtaatcACCAATATTTTCCAGCCATTGAAC | | |
| pBAD araC to pJRD ApaI fwd | attacgcgttaacccgggccGCCGTCAATTGTCTGATTCGTTA | Used to amplify fragments spanning from 27-bp upstream of the *araC* stop codon to 30-bp downstream of the *rrnB* T2 terminator in pBAD33.1F and derivatives. | Compatible for Gibson assembly into pJRD215 and derivatives linearized with ApaI. |
| pBAD rrnB to pJRD ApaI rev | atcatatgcatccgcgggccAAAATAAACAAAAGAGTTTGTAGAAACGCA | | |
| RK2 TcR_XhoI fwd | gcttggtcggtcatttcgccAGTCGCCTTGACCCGCAT | Used to amplify a TetA-TetR cassette from pCM62 to replace the KmR cassette in | Compatible for Gibson assembly into pJRD215 |

| | | | |
|---|---|---|---|
| RK2 TcR_BmtI-HF rev | cccccctgcagccaagctagTTAAGCCAGCCCCGACAC | pJRD215. | linearized with XhoI and BmtI-HF. |
| pBAD araC to pJRD BamHI fwd | ccccccccctgcaggtcgacgGCCGTCAATTGTCTGATTCGTTA | Used to amplify fragments spanning from 27-bp upstream of the *araC* stop codon to 30-bp downstream of the *rrnB* T2 terminator in pBAD33.1F and derivatives. | Compatible for Gibson assembly into pJRD215 and derivatives linearized with BamHI. |
| pBAD rrnB to pJRD BamHI rev | ttctacttatggtacccgggAAAATAAACAAAAGAGTTTGTAGAAACGCA | | |
| pirA Q f | TTAGCCACTTTCCAGCCGC | Primers for RT-qPCR amplification of *pirA/Bvp* and *gyrB* (an endogenous control gene). Reactions were performed using a Power SYBR Green RNA-to-CT 1-Step Kit (Applied Biosystems). | Primers described by Lin et al. (2022). |
| pirA Q r | CCGGAAGTCGGTCGTAGTGT | | |
| pirB Q f | TCGTTATCAGCCCACGCAG | | |
| pirB Q r | TTTCACCGATTCTGATGTGCA | | |
| gyrB Q 1 | GAAGGTGGTATTCAAGCGTTCG | | |
| gyrB Q 2 | GAGATGCCGTCTTCACGTTCT | | |

[a] Lowercase bases correspond to plasmid sequences, uppercase bases correspond to gene or flanking sequences

### Table S4 Alignment data for predicted effectors in D4

| *Vibrio parahaemolyticus* 13-306/D4 | | Literature homolog | | | Blastp results | | |
|---|---|---|---|---|---|---|---|
| Protein accession | Gene locus | Effector name | Protein accession | Source | Query coverage | E-value | % identity |
| WP_085609853.1 | ACMX29_RS08515 | VP1390 | BAC59653.1 | Jana et al. 2021 | 99% | 2E-154 | 26.93% |
| WP_024699427.1 | ACMX29_RS08525 | VP1388 | BAC59651.1 | Jana et al. 2021 | 97% | 0E+00 | 65.36% |
| WP_031852689.1 | ACMX29_RS08395 | VP1415 | BAC59678.1 | Salomon et al. 2014 | 63% | 2E-112 | 50.00% |
| WP_025789713.1 | ACMX29_RS24340 | TseVs | XDT02158.1 | Liu et al. 2025 | 79% | 3E-135 | 58.10% |
| WP_191786262.1 | ACMX29_RS07915 | T2Rhs-Nuc [BB22OP] | WP_015296737.1 | Tchelet et al. 2023 | 100% | 0E+00 | 95.19% |
| WP_020840162.1 | ACMX29_RS12530 | T2LipB [BB22OP] | WP_015296300.1 | Tchelet et al. 2023 | 100% | 0E+00 | 99.47% |
| WP_025550820.1 | ACMX29_RS06995 | T2Unk | WP_015313171.1 | Tchelet et al. 2023 | 100% | 3E-52 | 35.91% |
| WP_228767727.1 | ACMX29_RS20475 | T2Hydro [RIMD] | BAC61690.1 | Tchelet et al. 2023 | 100% | 0E+00 | 99.77% |
| WP_025542139.1 | ACMX29_RS13355 | Tme [T9109] | WP_047706523.1 | Fridman et al. 2020 | 62% | 4E-15 | 30.86% |
| WP_025520465.1 | ACMX29_RS18445 | Va16922 | WP_005376975.1 | Salomon et al. 2015 | 62% | 1E-11 | 27.20% |

Alignments were performed using the NCBI blastp suite set to default parameters. Protein sequences from NCBI assembly GCF_050888945.2 were used as queries against experimentally validated literature homologs.

### Table S5 Alignment data for predicted effectors in 297B

| *Vibrio parahaemolyticus* 12-297/B | | Literature homolog | | | Blastp results | | |
|---|---|---|---|---|---|---|---|
| Protein accession | Gene locus | Effector name | Protein accession | Source | Query coverage | E-value | % identity |
| WP_085577065.1 | B5C30_RS15275 | VP1390 | BAC59653.1 | Jana et al. 2021 | 100% | 0E+00 | 96.71% |
| WP_029793730.1 | B5C30_RS15265 | VP1388 | BAC59651.1 | Jana et al. 2021 | 100% | 0E+00 | 100.00% |
| WP_237771759.1 | B5C30_RS02625 | VP1415 | BAC59678.1 | Salomon et al. 2014 | 66% | 3E-31 | 30.10% |
| WP_029857615.1 | B5C30_RS14465 | PoNe | WP_029857615.1 | Jana et al. 2019 | 100% | 0E+00 | 100.00% |
| WP_025789713.1 | B5C30_RS04275 | TseVs | XDT02158.1 | Liu et al. 2025 | 79% | 3E-135 | 58.10% |
| WP_236575995.1 | B5C30_RS19400 | T2Rhs-Nuc [BB22OP] | WP_015296737.1 | Tchelet et al. 2023 | 91% | 0E+00 | 97.00% |
| WP_031856000.1 | B5C30_RS10045 | T2LipB [BB22OP] | WP_015296300.1 | Tchelet et al. 2023 | 100% | 0E+00 | 92.59% |

Alignments were performed using the NCBI blastp suite set to default parameters. Protein sequences from NCBI assembly GCF_002114585.1 were used as queries against experimentally validated literature homologs.

### Table S6 Additional comparisons of WP_025789713.1 to homologous proteins

| Homolog | | | Blastp Results | | | Genomic context | |
|---|---|---|---|---|---|---|---|
| Species and strain | Protein accession | Annotation | Query coverage | E-value | % identity | Upstream feature | Downstream feature |
| *Photobacterium damselae* A-162 | WP_069531219.1 | hypothetical protein | 78% | 4.0E-175 | 71.1% | Type VI secretion system VgrG | hypothetical protein |
| *Aliivibrio fischeri* ZF47 | WP_017018897.1 | hypothetical protein | 74% | 1.0E-144 | 57.7% | DUF4123 domain-containing protein | hypothetical protein |
| *Vibrio vulnificus* VA-WGS-18041 | WP_130243383.1 | hypothetical protein (predicted RIX motif) | 75% | 4.0E-136 | 56.1% | hypothetical protein | hypothetical protein |
| *Vibrio amylolyticus* ZSDE26 | WP_248008649.1 | DUF4150 domain-containing protein | 73% | 2.0E-138 | 61.1% | Type VI secretion system DotU | hypothetical protein |
| *Salinivibrio proteolyticus* YCSC6 | WP_346426280.1 | MIX_V domain-containing protein | 72% | 9.0E-130 | 59.0% | BioA aminotransferase | hypothetical protein |
| *Pantoea* sp. At-9b | ADU69816.1 | hypothetical protein | 57% | 7.0E-14 | 22.0% | Type VI secretion system Hcp | hypothetical protein |

Alignments were performed using the NCBI blastp suite set to default parameters. The protein sequence WP_025789713.1 was used as a query against the indicated homolog proteins.

### Table S7 Alignment data for predicted antibiotic resistance determinants in D4

| ***Vibrio parahaemolyticus* 13-306/D4** | | | Literature homolog | | | Blastp results | | |
|---|---|---|---|---|---|---|---|---|
| Protein accession | Gene locus | Gene product | Name | Protein accession | Source | Query coverage | E-value | % identity |
| WP_005463392.1 | ACMX29_RS16880 | Diacylglycerol kinase | DgkA | WP_005463392.1 | Saad et al. 2025 | 100% | 0E+00 | 100.00% |
| WP_025546543.1 | ACMX29_RS16885 | Phosphoethanolamine transferase | EptA | WP_005477155.1 | Saad et al. 2025 | 100% | 0E+00 | 99.64% |
| WP_031944061.1 | ACMX29_RS12335 | Tetracycline resistance MFS efflux pump | Tet(B) | WP_031944061.1 | Han et al. 2015 | 100% | 0E+00 | 100.00% |
| WP_000088605.1 | ACMX29_RS12330 | Tetracycline resistance regulatory protein | TetR | WP_000088605.1 | Han et al. 2015 | 100% | 0E+00 | 100.00% |
| WP_005459715.1 | ACMX29_RS21045 | CARB family class A beta-lactamase | VPA0477 | WP_005479236.1 | Chiou et al. 2015 | 100% | 0E+00 | 99.29% |

Alignments were performed using the NCBI blastp suite set to default parameters. Protein sequences from NCBI assembly GCF_050888945.2 were used as queries against antibiotic resistance determinants previously reported in the literature.

#### Table S8 Relative expression of *pirA/B*$^{vp}$ compared to D4 wild type

| Gene | Strain | Mean $\Delta C_q$ | $\Delta\Delta C_q$ | Fold change ($2^{-\Delta\Delta C_q}$) | 95% CI | Adjusted *P* (Dunnett) |
|---|---|---|---|---|---|---|
| *pirA vp* | D4 wild type | -6.33 | 0.00 | 1.00 | — | — |
| *pirA vp* | D4 Δ*hcp1* | -10.05 | -3.72 | 13.18 | 1.1–160 | 0.0436 |
| *pirA vp* | D4 Δ*hcp2* | -7.35 | -1.02 | 2.03 | 0.17–25 | 0.7574 |
| *pirA vp* | D4 Δ*hcp1/2* | -8.92 | -2.60 | 6.05 | 0.50–74 | 0.1642 |
| *pirB vp* | D4 wild type | -7.85 | 0.00 | 1.00 | — | — |
| *pirB vp* | D4 Δ*hcp1* | -11.61 | -3.76 | 13.52 | 1.2–157 | 0.0383 |
| *pirB vp* | D4 Δ*hcp2* | -8.82 | -0.97 | 1.96 | 0.17–23 | 0.7740 |
| *pirB vp* | D4 Δ*hcp1/2* | -10.40 | -2.55 | 5.84 | 0.50–68 | 0.1642 |

Values indicate fold change relative to D4 wild type ($2^{-\Delta\Delta C_q}$), normalized to *gyrB*; $\Delta C_q = C_q(target) - C_q(gyrB)$, and $\Delta\Delta C_q$ = mean $\Delta C_q$(mutant) − mean $\Delta C_q$(wild type). 95% CIs and adjusted *P* values are from Dunnett's multiple comparisons test on $\Delta C_q$ values (n = 3 biological replicates per strain).

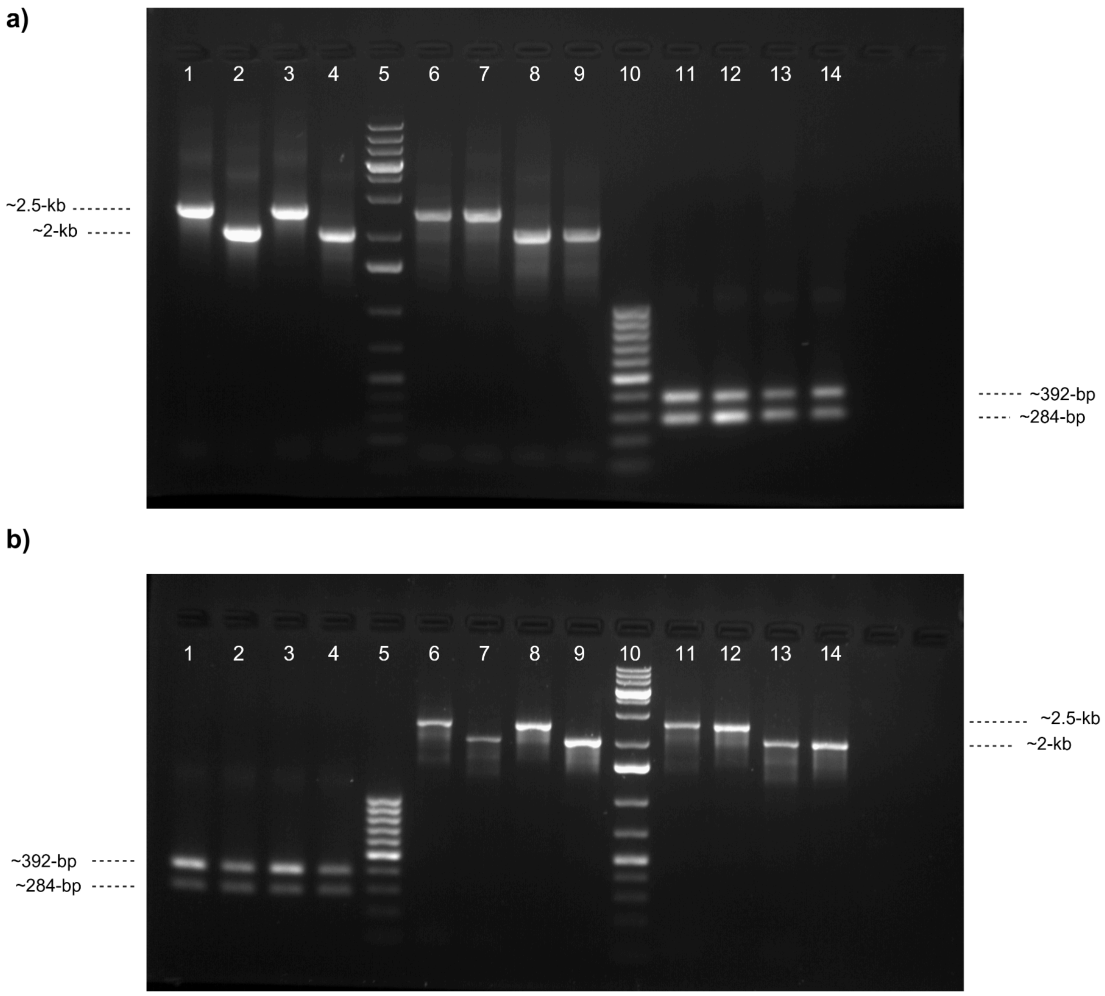


**Fig. S1 Agarose gels from PCR validation of T6SS knockout mutants of D4 (a) and 297B (b).** Mutants obtained by allelic exchange were verified using primers that anneal 1kb upstream and 1kb downstream of the *hcp1* (519bp) or *hcp2* (480bp) gene and a genomic DNA template from the indicated strain. A duplex PCR screen was also performed to confirm retention of the *pirA*$^{vp}$ and *pirB*$^{vp}$ genes. Samples in (**a**) are as follows:

Lanes 1-4, *hcp1* screens of (1) D4 wt, (2) D4 Δ*hcp1*, (3) D4 Δ*hcp2*, and (4) D4 Δ*hcp1/2*; Lane 5, GeneRuler 1kb+ ladder; Lanes 6-9, *hcp2* screens of (6) D4 wt, (7) D4 Δ*hcp1*, (8) D4 Δ*hcp2*, and (9) D4 Δ*hcp1/2*; Lane 10, GeneRuler 100bp ladder; Lanes 11-14, *pirA/B*$^{vp}$ duplex PCR screens of (11) D4 wt, (12) D4 Δ*hcp1*, (13) D4 Δ*hcp2*, and (14) D4 Δ*hcp1/2*

Samples in (**b**):

Lanes 1-4, *pirA/B*$^{vp}$ duplex PCR screens of (1) 297B wt, (2) 297B Δ*hcp1*, (3) 297B Δ*hcp2*, and (4) 297B Δ*hcp1/2*; Lane 5, GeneRuler 100bp ladder; Lanes 6-9, *hcp1* screens of (6) 297B wt, (7) 297B Δ*hcp1*, (8) 297B Δ*hcp2*, and (9) 297B Δ*hcp1/2*; Lane 10, GeneRuler 1kb+ ladder; Lanes 11-14, *hcp2* screens of (11) 297B wt, (12) 297B Δ*hcp1*, (13) 297B Δ*hcp2*, and (14) 297B Δ*hcp1/2*

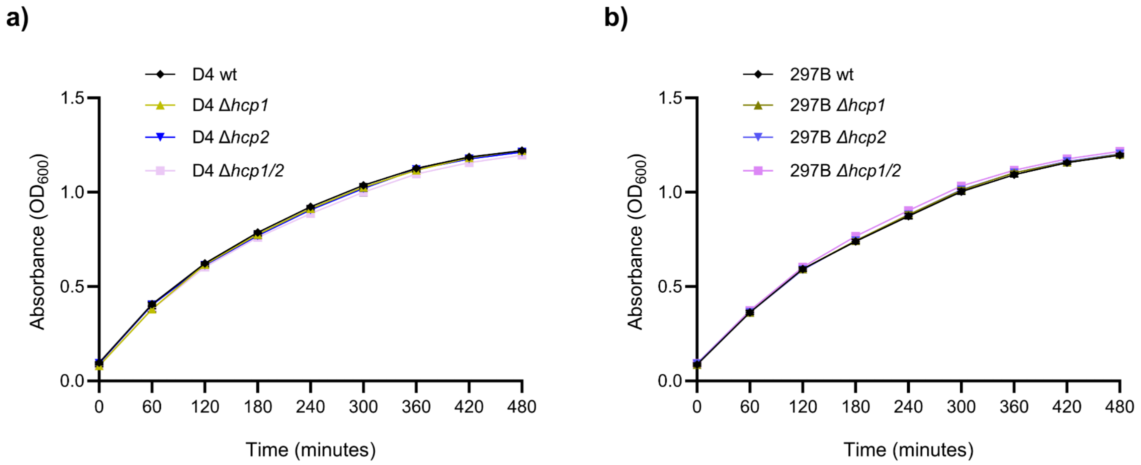


**Fig. S2 Growth over time of wild type and mutant strains of D4 (a) and 297B (b), measured by absorbance at 600 nm ($OD_{600}$).** Overnight cultures of wild type and mutant strains were normalized to an OD600 of 0.1 in fresh MLB and aliquoted (200 µL per well) into a 96-well plate. Plates were incubated in a BioTek Cytation 5 microplate reader at 30°C with constant shaking. Data shown represent mean values from three independent experiments.

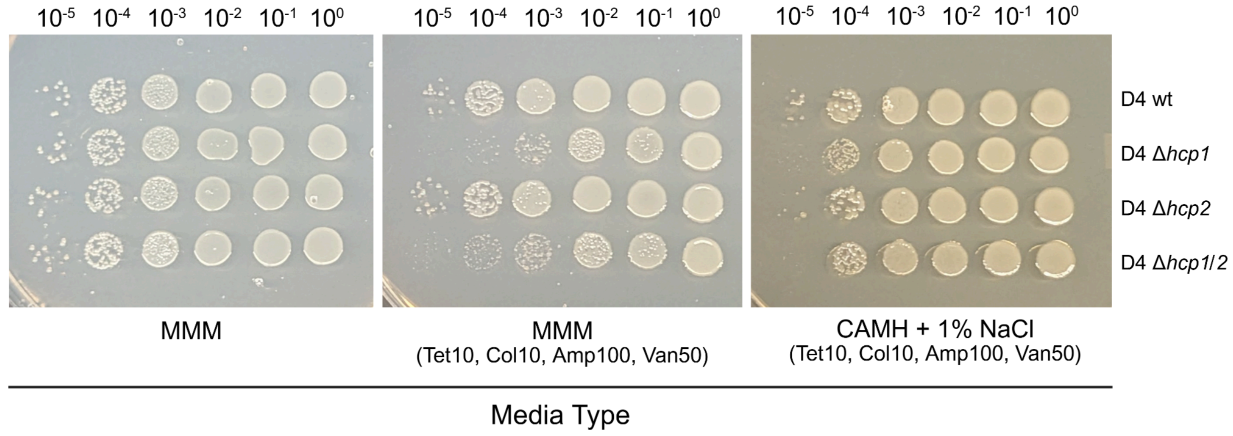


**Fig. S3 Growth of D4 wild type and mutant strains on antibiotic-supplemented media plates.** Overnight cultures of the indicated D4 strains were prepared in CAMH + 1% NaCl at 30°C with constant shaking. The following day, cultures were normalized to a concentration of approximately 1.5E8 CFU/mL. Serial 10-fold dilutions were then spotted onto MMM and CAMH + 1% NaCl agar plates supplemented with 10 µg/mL tetracycline (Tet10), 10 µg/mL colistin (Col10), 100 µg/mL ampicillin (Amp100), and 50 µg/mL vancomycin (Van50), or onto unsupplemented MMM. Plates were briefly allowed to dry, and incubated at 30°C for 24h before inspection.
